# Pharmacological targeting of SOS1-RAS interaction triggers pancreatic β-cell proliferation and sustainably reverses diabetic hyperglycemia

**DOI:** 10.1101/2025.06.03.657628

**Authors:** Adriana Papadimitropoulou, Chrysanthi Charalampous, Paraskevi Kogionou, Diana Reinhardt, Johanna Sonntag, Anthony Gavalas, Marco H. Hofmann, Patrik Erlmann, Michael Franti, Jonas Doerr, Thomas Klein, Gareth R. Willis, Ioannis Serafimidis

## Abstract

Clinical studies have suggested that restoring a sufficient mass of functional β cells can provide an effective treatment option for diabetes, however, it remains unclear whether this can be achieved through pharmacological stimulation of endogenous β-cell proliferation. We demonstrate here that ectopic expression of a constitutively active form of Kras (Kras^G12D^) exclusively in pancreatic endocrine cells suppresses β-cell proliferation, resulting in a dramatic reduction in β-cell numbers and islet size. Conversely, we demonstrate that the potent and selective SOS1-RAS interaction inhibitor BI-3406 promotes unprecedented levels of β-cell proliferation in primary human islets, both in culture and following transplantation in immunocompromised diabetic mice. Importantly, using murine models of streptozotocin-induced diabetes, we show that BI-3406 treatment restores β-cell mass, leading to a gradual normalization of blood glucose and insulin levels, as well as to sustainable improvement in glucose tolerance. Our data provide the first pre-clinical evidence of an orally bioavailable KRAS inhibitor that can directly induce β-cell regeneration.

## INTRODUCTION

Diabetes mellitus is a chronic metabolic disorder characterized by elevated blood glucose levels resulting from insufficient insulin production and/or impaired insulin action. Both type 1 and type 2 diabetes involve a decline in β-cell mass^1,2^, the primary source of insulin production in the pancreas. Restoring a functional β-cell mass represents a promising approach for treating diabetes, and it has been proposed that induction of islet regeneration through *in vivo* stimulation of β-cell proliferation could be an attractive strategy for achieving long-term glycemic control^3–5^.

Over the years, extensive research has shed light on the possibility of inducing adult human β-cells to proliferate^6^, challenging the previous notion that β-cell proliferation is unattainable after the early neonatal period. Unlike many other tissues in the body, the regenerative capacity of adult pancreatic islet cells is only sufficient to maintain a relatively stable endocrine cell mass, but unable to compensate for significant β-cell loss^7,8^. Nevertheless, adult β-cells maintain the capacity to achieve high-rate cell proliferation when the right stimulus is applied, offering hope that β-cell mass restoration may be a viable option after diabetic loss^6^.

Considerable progress has been made in understanding β-cell proliferation under physiological conditions and in pathology. Studies in mice, particularly during development, have elucidated many molecular components essential for β-cell proliferation^9,10^. For example, a wide range of secreted factors, including GLP-1^11,12^, Adipsin^13^, SerpinB1^14^, IGF-1^15^, IGF-2^16^, HGF-1^17^ and others, have been shown to stimulate β-cell proliferation in dynamic physiological conditions. Additionally, extracellular matrix components also play a critical role in creating an islet microenvironment that supports the proliferative capacity of β-cells through localized growth factor signalling^18^. Despite accumulating knowledge about extracellular signals that stimulate β-cell proliferation, the challenge now lies in identifying the signaling pathways that transduce these signals within β-cells. The most successful attempt to identify a mitogenic pathway in β-cells has been the implication of the Calcineurin/NFAT/DYRK1A pathway in this process^19^, and the demonstration that DYRK1A inhibitors, particularly Harmine, can stimulate adult human β-cell proliferation to a considerable extent^20–22^. The success of Harmine as a potent preclinical β-cell proliferative factor and the subsequent controversy on its clinical safety as a monotherapy due to possible neurotoxicity, cardiovascular and other side-effects (discussed in^6^), prompted the search for additional candidates targeting other classes of mitogenic pathways. In this context, stimulating the MAP kinase pathway or inhibiting the CREB^23^, GSK3^14^, TGFβ^24^ or KRAS^25^ signaling pathways have all emerged as attractive targets to enhance β-cell proliferation.

KRAS, a member of the RAS superfamily of small GTPases, acts as a molecular switch that transduces signals from cell surface receptors to downstream effectors, ultimately modulating a multitude of diverse and often opposing cellular processes such as proliferation, differentiation, survival and apoptosis^26,27^. The case of the pancreas is a particularly striking example of this diversity of actions: whereas activating KRAS mutations stimulate transformation and tumor formation in the exocrine compartment, they never cause pancreatic endocrine tumors^28^. Conversely, endocrine tumors, including insulinomas, are never associated with oncogenic mutations in KRAS, suggesting that cell proliferation within the endocrine compartment is regulated differently^29,30^.

Genetically engineered mouse models have offered valuable insights into the role of KRAS signaling in pancreatic β-cells. For example, KRAS happloinsufficiency through the genetic ablation of one Kras allele in β-cells leads to a remarkable increase in β-cell neogenesis in the fetal pancreas, followed by rapid β-cell expansion during the perinatal period and significantly improved glucose tolerance throughout adult life^25^. Conversely, the conditional activation of Kras^G12D^, a constitutively active mutant form of KRAS, in the entire pancreatic epithelium leads to a notable reduction in β-cell proliferation and subsequent β-cell mass decline^25^. Importantly, these mice also exhibit hyperglycemia, indicating a direct connection between aberrant KRAS signaling and the development of a diabetic phenotype^25^. We provide here additional evidence in support of these observations, by generating a mouse model where Kras^G12D^ expression is restricted to pancreatic endocrine progenitor cells and their progeny, allowing an assessment of the direct and cell-autonomous impact of constitutively active KRAS on β-cell development and function. In these mice, we observed a marked reduction in islet numbers and a significant decline in both insulin-producing β-cells and glucagon-producing α-cells, concomitantly with the development of severe hyperglycemia, highlighting the role of KRAS signaling as a negative regulator of β-cell growth.

A model was previously proposed in which KRAS activates multiple and opposing downstream effectors in endocrine cells, including the pro-proliferative MAPK pathway and the antagonistic anti-proliferative tumour-suppressor RASSF1A pathway^25,31^. The dominance of the antiproliferative effect of KRAS in β cells was found to depend on the expression of another tumour suppressor, menin^32^, which prevents the MAPK pathway from driving proliferation but does not affect inhibitory pathways like RASSF1A. Loss of menin leads to β-cell proliferation due to the removal of the MAPK-driven proliferation blockage, while loss of KRAS signaling increases proliferation by reducing RASSF1A activity^31^. This model therefore predicts that pharmacological inhibition of KRAS may be a promising therapeutic strategy to stimulate β-cell proliferation, however, until recently, KRAS was considered an “undruggable” target due to its small size and smooth and shallow surface that is refractory to small molecule binding^33,34^. Hofmann et al.^35^ recently reported the discovery of a highly potent, selective, and orally bioavailable small molecule, BI-3406, that binds to the catalytic domain of the RAS partner SOS1, thereby preventing their interaction and downstream signalling^35^. BI-3406 therefore acts as an all-purpose pan-H, N and KRAS inhibitor with a demonstrated ability to limit cellular proliferation in a broad range of KRAS-driven cancers, sparking a renewed interest on the clinical use of KRAS inhibition in anti-cancer treatment, as well as towards novel disease indications^35,36^.

Our study aims to comprehensively investigate the implications of KRAS signaling attenuation in β-cell proliferation, with a specific focus on the therapeutic potential of BI-3406. Using well-characterized mouse models of chemically-induced diabetes^37^, we demonstrate that KRAS inhibition gradually reverses hyperglycemia, restores islet morphology, and improves β-cell functionality. In addition to the in vivo studies, we explore the effects of KRAS inhibition on primary human islets, revealing a remarkable potency in stimulating β-cell proliferation both in culture and when transplanted in immunocompromised diabetic mice.

In summary, our results highlight the potential of targeting KRAS signaling via SOS1 inhibition, as a means to induce β-cell regeneration and restore insulin production in individuals with diabetes.

## RESULTS

### Kras activity suppresses growth of pancreatic endocrine cells

We previously described the Tg^Pdx1Cre^-Actb^LSLKras*^ (Pdx1Cre-AKras*) mouse model (Fig S1A), which rapidly develops Pancreatic Ductal Adenocarcinoma (PDAC) in a stagewise and orderly fashion that faithfully simulates the human condition^38,39^. Interestingly, in addition to the progressive development of invasive PDAC, we observed that Pdx1Cre-AKras* mice also developed hyperglycemia, as indicated by the consistently higher fasting blood glucose levels (average 1.53-fold increase) compared to age-matched controls (Fig. S1B). To investigate whether the observed hyperglycemia is due to Kras overactivation specifically on pancreatic endocrine cells and not a secondary complication of PDAC development, we generated the Tg^Ngn3Cre^-Actb^LSLKras*^ (Ngn3Cre-AKras*) mouse model (Fig 1A). In this model, the mutated and constitutively active form of Kras (Kras^G12D^ or Kras*) is expressed under the control of the strong β-actin promoter, exclusively in the endocrine progenitor cells and their progeny. To validate the specificity of the Ngn3-Cre driver, we first generated Tg^Ngn3Cre^-Rosa26^LmTdTL-mEGFP^ mice (Ngn3Cre-RosaGFP), where the expression of Cre recombinase permanently labels target cells with mEGFP (Fig S1C). We observed a strong colocalization of mEGFP with endogenous Ngn3 expression at E14.5 (Fig S1D), and mEGFP+ cells were concentrated in the trunk of the developing pancreatic epithelium at E16.5, where endocrine progenitors reside (Fig S1D). Importantly, all islet cells in the adult transgenic mice were strongly positive for mEGFP, successfully marking the entire endocrine compartment as previously reported^40^ (Fig S1D).

**Figure 1.**
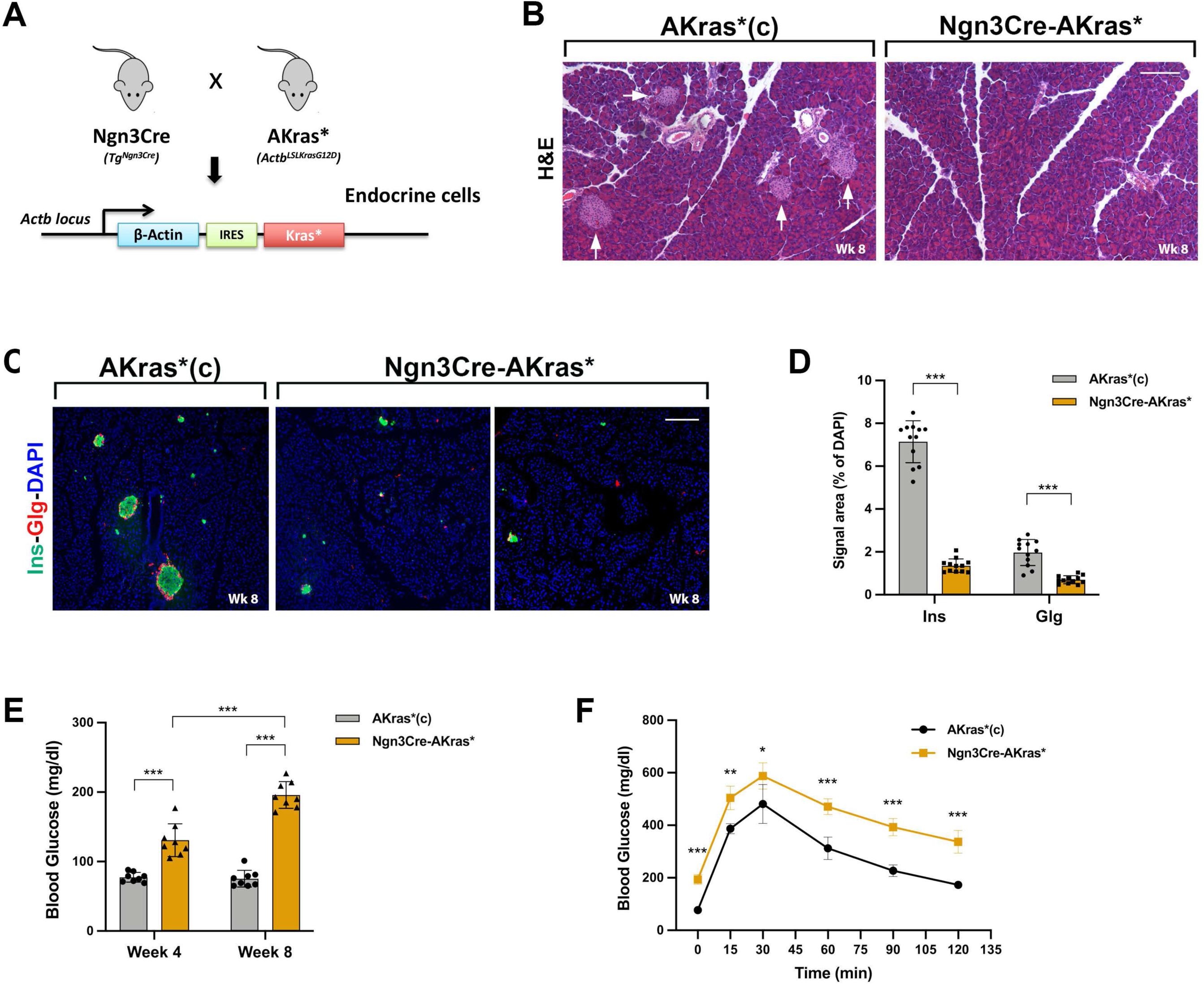
KRAS overactivation suppresses pancreatic islet growth. A) Ngn3Cre-AKras* mice that express a mutated (KRAS^G12D^) and constitutively active form of KRAS, under the control of the β-actin promoter, exclusively in the endocrine progenitor (Ngn3^+^) cells and their progeny, were analysed and compared to age-matched unrecombined AKras* controls. B) Representative images of the histological analysis of pancreata from adult (8-week-old) Ngn3Cre-AKras* mice (n=3) revealed no signs of PDAC, but a significant reduction in the number of islets (white arrows), compared to controls (n=3). C) Representative images of Insulin (Ins) and Glucagon (Glg) immunostainings from 8-week-old Ngn3Cre-AKras* mice (n=3) and AKras* controls (n=3), and quantitative analysis of Ins and Glg expression from these images (D) confirm the reduction in pancreatic islet mass. E) Fasting blood glucose levels at weeks 4 and 8 show a gradual induction of hyperglycemia in Ngn3Cre-AKras* mice (n=8) when compared to age-matched AKras* controls (n=8). F) Consequently, glucose uptake is significantly less efficient in 8-week-old Ngn3Cre-AKras* mice (n=5) compared to controls (n=5), as indicated by IPGT tests. All values are expressed as mean +/-SD. Data points in (D) represent measurements of individual pancreatic sections per genotype. Data were statistically analysed by unpaired two-tailed t tests. \**p*<0.05, \*\**p*<0.01 and \*\*\**p*<0.001. Scale bars, 120 μm.

Ngn3Cre-AKras* adult mice did not exhibit signs of pancreatic cancer (PDAC), but we observed a significant reduction in the number of islets (Fig 1B). More specifically, immunofluorescent analysis and islet-cell quantitation revealed a 5.26-fold reduction in Ins+ β-cells and a 2.82-fold reduction in Glg+ α-cells when Kras was continuously expressed and constitutively active in these cells (Fig 1C, D). The loss of islet cells correlated with a progressive induction of severe hyperglycemia, as evidenced by the measurement of fasting blood glucose levels (Fig 1E). At 4 weeks of age, Ngn3Cre-AKras* animals exhibited an average blood glucose concentration of 131 (+/-24) mg/dl (1.69-fold higher compared to controls), which increased to 196 (+/-19) mg/dl by 8 weeks of age (2.61-fold higher compared to control mice) (Fig 1E). Consequently, an IPGT test on 8-week-old Ngn3Cre-AKras* mice demonstrated that these were significantly less efficient in clearing excess glucose from circulation compared to age-matched controls (Fig 1F), further confirming their diabetic phenotype.

Our results provide crucial genetic evidence that Kras activity suppresses β-cell growth in a cell-autonomous manner, therefore suggesting that targeted inhibition of Kras signaling can potentially exert the opposite effect.

### SOS1-RAS inhibitor BI-3406 induces β-cell proliferation in human islets

We examined the effects of the SOS1-RAS inhibitor BI-3406 on intact human pancreatic islets from 6 donors aged 18-63 (Fig 2A, B). The optimum dosage for BI-3406 in cultured human islets was determined at 300nM (Fig S2A, B; details in Methods).

**Figure 2.**
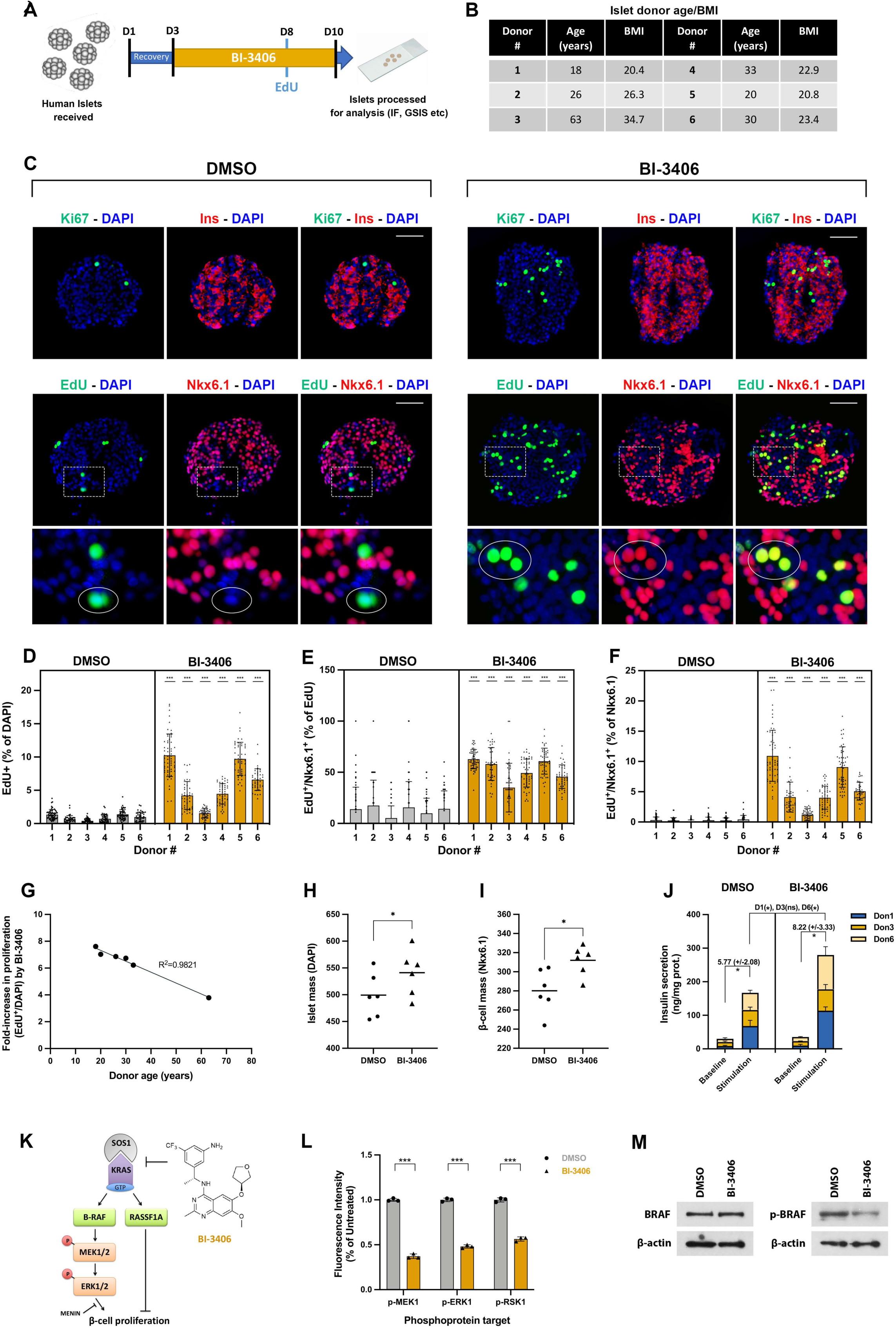
BI-3406 induces β-cell proliferation in human islets. A) Schematic timeline illustrating the course of treatment of human islets with BI-3406 (D: Day of treatment). B) List of human islet samples used in this study, including the respective donor’s age and BMI. C) Representative images of islet sections from Donor 1 samples following treatment with BI-3406 or DMSO, immunostained for Insulin (Ins), Ki67 and Nkx6.1, as well as for the fluorescent detection of EdU incorporation. Bottom row panel exhibits high-magnification views of regions denoted by dashed squares, showcasing cells double-labelled with EdU and Nkx6.1. (D-F) Quantitative analysis of the impact of BI-3406 treatment on islet cell proliferation for each donor: D) Quantitation of EdU^+^ cells, expressed as a percentage of islet DAPI; E) Quantitation of EdU/Nkx6.1 double-labelled cells, expressed as a percentage of EdU^+^ cells; F) Quantitation of EdU/Nkx6.1 double-labelled cells, expressed as a percentage of Nkx6.1^+^ cells. G) Plot illustrating the fold-increase in proliferative (EdU+) cells in response to BI-3406 for each donor, relative to their age, revealing a linear correlation with an R^2^ value of 0.9821. H) BI-3406 treatment increases islet mass, expressed as the total number of DAPI^+^ nuclei per islet section. H) Similarly, BI-3406 treatment increases β-cell mass, expressed as the total number of Nkx6.1^+^ cells per islet section. J) Improved glucose-stimulated insulin secretion (GSIS) was observed in BI-3406 treated islets, particularly in younger donors (Donors 1 and 6): an 8.0-, 3.9- and 5.3-fold increase in insulin secretion between baseline and stimulation conditions was recorded for untreated Donor 1, 3 and 6 islets respectively. These values rose to 11.4-, 4.8- and 8.5-fold for BI3406-treated islets of the same donors. K) Schematic model of the SOS1-KRAS signaling pathway, indicating downstream effectors and their role in regulating β-cell proliferation. L) Luminex assays for key phosphoprotein targets of signaling pathway involved in cell proliferation, demonstrated a 2.69-, 2.08, and 1.77-fold reduction in the expression levels of the MAPK components p-MEK1, p-ERK1 and p-RSK1 respectively, in lysates of Donor 6 human islets treated with BI-3406. M) Western blot for BRAF and phospho-BRAF on Donor 6 islet lysates also revealed a reduction in the expression levels of the phosphorylated form of BRAF in response to BI-3406. All values are expressed as mean +/-SD. Data points in (D-E) represent measurements of individual islets per donor, per condition (n=58 for Donor 1; n=40 for Donor 2; n=50 for Donors 3-5; n=40 for Donor 6). Data points with zero value are not visible on the x-axis of (D-F). Data points in (G-I) represent average values per donor, per condition. Data were statistically analysed by paired (J; Baseline vs Stimulation) or unpaired (D-F, H-J, L) two-tailed t tests. \**p*<0.05, \*\**p*<0.01 and \*\*\**p*<0.001. Scale bars, 40 μm.

We initially assessed islet-cell proliferation by measuring EdU labelling and Ki67 expression via immunostaining. Our immunofluorescent analysis demonstrated a significant increase in the number of Ki67+ and EdU+ cells in BI-3406-treated islets (Fig 2C). We quantified EdU labelling on each of the six donor samples and found an average 6.38-fold (+/-1.34) increase in the number of EdU+ cells in BI-3406-treated islets compared to controls (Fig 2D). Importantly, the percentage of EdU+ cells co-expressing the mature β-cell marker Nkx6.1 increased on average from 12.77% (+/-4.43) in control islets to 51.97% (+/-10.67) in BI-3406-treated islets (Fig 2E), indicating that BI-3406 was particularly effective in inducing β-cell-specific proliferation (Fig. 2C; bottom panel). Conversely, the percentage of Nkx6.1+ cells simultaneously labelled with EdU increased by a striking 19.98-fold (+/-10.54) in BI3406-treated islets (Fig 2F). For comparison, the percentage of Glucagon+ α-cells simultaneously labelled with EdU was found increased by just 1.99-fold in BI-3406-treated islets (Fig S2C, D).

Interestingly, we observed a strong linear correlation between the fold-increase in EdU+ islet cells and the age of the islet donors, with younger islets exhibiting up to 2.5-fold higher increase in proliferation indices compared to older ones (Fig 2G). An analogous correlation was observed between BI-3406-induced proliferation and islet donor body-mass index (BMI) (Fig S2E). A direct comparison of the activity profiles of BI-3406 and Harmine, assessed in parallel and under identical conditions, demonstrated that BI-3406 consistently performed better than Harmine, and was on average 30.1% (+/-4.88) more effective in stimulating β-cell proliferation (Fig S2F).

Next, we asked whether the observed increase in islet proliferation marker expression in response to BI-3406 translates to a similar increase in islet cell numbers and functionality. We found that total islet mass was increased by 8.42% (+/-3.36%) (Fig 2H), while β-cell mass was increased by 11.66% (+/-5.46%) in BI-3406-treated islets compared to controls (Fig 2I). We also performed static glucose-stimulated insulin secretion assays (GSIS) on BI-3406-treated islets, which on average displayed a higher fold-stimulation of insulin release compared to controls, particularly evident in islets from younger donors (Fig 2J).

To determine whether KRAS signalling is attenuated in human islets treated with BI-3406, we evaluated expression of key targets of the KRAS signaling cascade (Fig 2K). Molecular profiling of treated and control islets using a panel of 20 signaling phosphoproteins in a Luminex-based biomarker screening assay, indicated that expression of the MAPK signaling components MEK1, ERK1, and RSK1 were significantly decreased upon treatment with BI-3406 (Fig 2L). Moreover, we demonstrated that the activated (phospho-S445) form of the immediate downstream KRAS effector, pBRAF, was also significantly decreased in BI3406-treated islets, providing further support to the evidence that KRAS signaling is attenuated in response to BI-3406 (Fig 2M). Interestingly, all other target phosphoproteins in the Luminex panel were either unaffected by the treatment, or non-detectable in the protein lysates, with the notable exception of STAT3 that appeared to be induced by BI-3406 (Fig S2G; see Discussion).

In conclusion, our results demonstrate a strong age-dependent induction of β-cell proliferation in human islets treated with the SOS1-RAS inhibitor BI-3406, leading to larger islets with improved β-cell functionality *in vitro*.

### BI-3406 increases β-cell mass and reverses hyperglycemia in diabetic mice

Next, we evaluated the impact of BI-3406 on diabetic mice in vivo. Chemical diabetes was induced on male c57/Bl6 mice using the “multiple, low dose” streptozotocin (STZ) administration scheme^37^, as described in Methods. This scheme is characterized by a delayed onset of hyperglycemia and avoids complete damage of pancreatic islets, preserving a minimal mass of β-cells^37^. We administered BI-3406 at 50 mg/kg orally, twice per day (with a 6h interval), for 30 consecutive days (Fig 3A). This administration scheme was well tolerated by the diabetic mice, which remained active and agile throughout the course of treatment. A modest reduction in body weight was observed upon the initiation of BI-3406 administration, primarily attributed to a temporary decrease in appetite (Fig S3A). However, this decrease was completely reversed, and body weight fully regained, by the end of the treatment period (Fig S3A).

**Figure 3.**
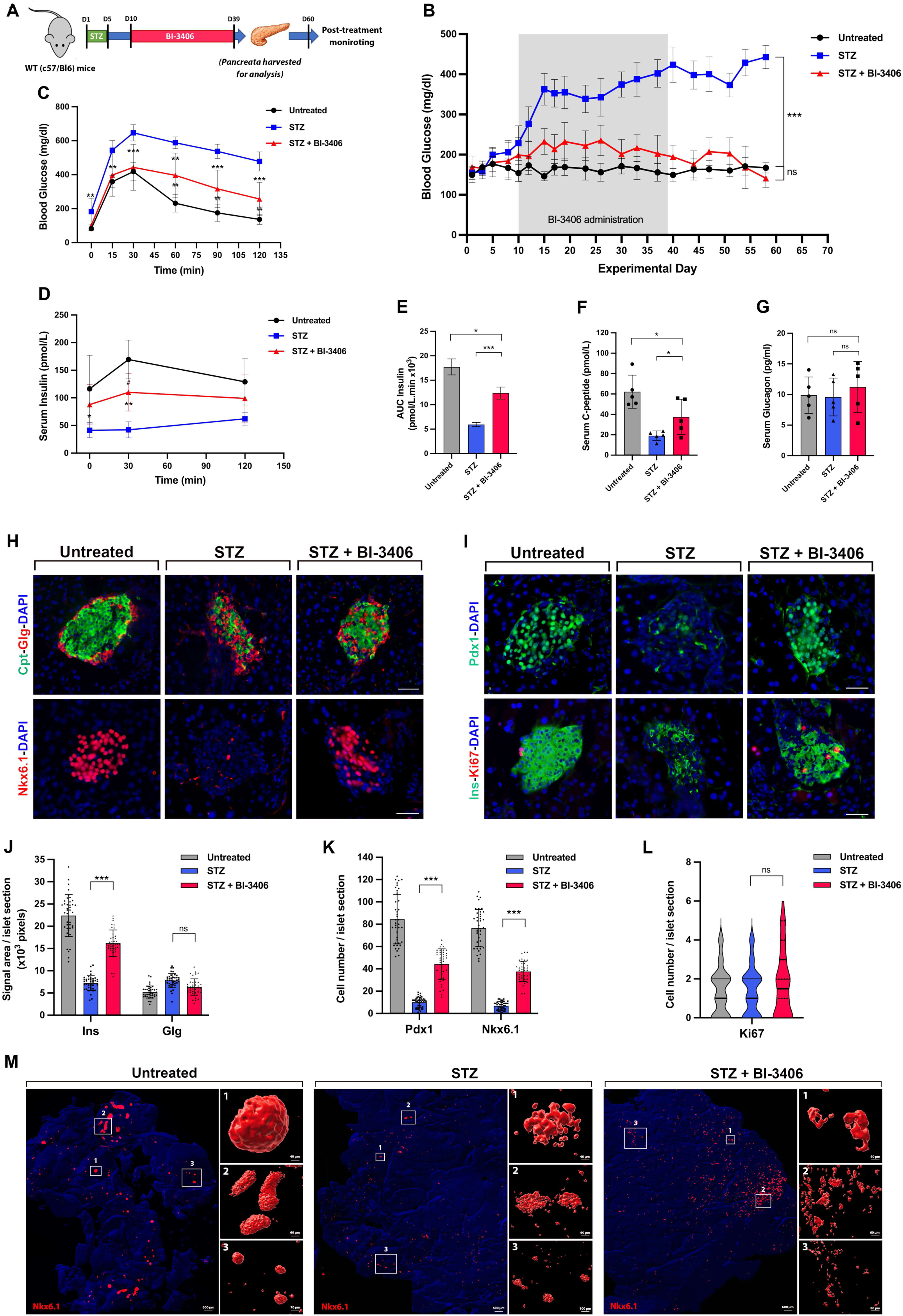
BI-3406 increases β-cell mass and reverses hyperglycemia in diabetic mice. A) Schematic representation of the *in vivo* treatment timeline, including schedule of STZ and BI-3406 administration. B) Monitoring of blood glucose levels throughout treatment revealed that diabetic mice receiving BI-3406 (STZ+BI-3406; n=14) maintained lower glucose levels compared to diabetic controls (STZ; n=16), eventually reaching normoglycemic levels similar to untreated controls (n=17). C) IPGT tests conducted on mice from all three experimental groups at the end of treatment (D40), demonstrated significantly improved blood glucose clearence in BI-3406-treated mice (n=10) compared to STZ controls (n=10), without, however, reaching the rates of untreated controls (n=13). D-E) An ELISA for insulin measured during IPGTT at D40 (D), and the corresponding calculations of the areas under the curve (AUC) (E), showed that total serum insulin levels in the BI-3406-treated group were significantly higher compared to STZ control mice, but did not reach the levels observed in untreated controls. F) An ELISA for c-peptide measured at D40 closely mirrored the results obtained for insulin. G) An ELISA for glucagon measured at D40 revealed no significant changes in serum glucagon levels between the three groups. (H-I) Representative images of pancreatic islet sections from each experimental group at D40 (n=3 per group), immunostained for C-peptide, Glucagon and Nkx6.1 (H), as well as Insulin, Pdx1 and Ki67 (I), demonstrate a significant restoration of islet mass in BI-3406-treated mice compared to STZ diabetic controls. J) Quantitative assessment of Insulin and Glucagon from the corresponding immunofluorescent images revealed a significant restoration of β-cell numbers in response to BI-3406, but no significant effect on α-cells. K) Quantitative assessment of Pdx1 and Nkx6.1 from the corresponding immunofluorescent images confirmed a partial restoration of mature and functional β-cells upon treatment with BI-3406. L) Quantitative assessment of Ki67 from the corresponding immunofluorescent images demonstrated no significant proliferation by BI-3406 at the time of harvest (D40). M) FluoClearBABB (iDisco) images of whole pancreatic fragments from the three experimental groups, immunostained for Nkx6.1 to depict the distribution of whole islets in the 3D pancreatic space. Selected areas (marked by white squares) were surface-rendered (details in Methods) and presented in high magnification on the right of the corresponding main images. FluoClearBABB imaging demonstrated that the restored β-cells in the BI-3406-treated samples did not reorganize in clusters and did not acquire a wild-type islet morphology. Values in (B-D, F-G, J-L) are expressed as mean +/-SD, and in (E) as mean +/-SEM. Data points in (B, C) represent average values per group. Data points in (D, F, G) represent averages from triplicate ELISA measurements per serum sample (n=5 mice per group). Data points in (J-L) represent measurements of individual islet sections per condition (n=40 per group). Data points with zero value are not visible on the x-axis of (L). Data were statistically analysed by one-way ANOVA with Tukey’s post-hoc test. \**p*<0.05, \*\**p*<0.01, \*\*\**p*<0.001 and ns (not significant). Scale bars: (H-I) 40 μm, (M) as indicated on each image.

We monitored blood glucose levels in untreated-controls, STZ-controls, and STZ+BI3406-treated mice, twice a week following a 4h fasting, for the entire course of the study (Fig 3B). Untreated mice (100% of n=17) maintained steady blood glucose levels in the range of 120-180 mg/dl throughout the study. In contrast, STZ-control mice (94% of n=17) had a gradual increase in blood glucose levels, reaching steady hyperglycemia (320-420 mg/dl) from D15 onwards. Remarkably, the majority of BI3406-treated mice (82% of n=17) never reached hyperglycemia and only had a small increase in blood glucose levels (up to 235 mg/dl), which regressed to nearly WT levels (140-200 mg/dl) and remained steadily low even two weeks after the last BI-3406 dose administration (Fig 3B). Blood glucose levels for individual mice across the three experimental groups are presented in Fig S3B-D. WT animals receiving the same treatment with BI-3406 had no significant difference in their blood glucose levels compared to their untreated WT counterparts (Fig S3E).

We performed IPGT tests in mice from the three experimental groups at the end of treatment (D40; Fig 3C) as well as at the end of the post-treatment monitoring period (D60; Fig S3F). In both instances, BI-3406-treated mice were significantly more efficient in clearing excess glucose from their circulation compared to STZ-control mice. However, the rate of glucose clearance in the STZ+BI-3406 group, from 30’ onwards, was significantly lower than that of untreated controls, indicating that the nearly fully-restored fasting blood glucose levels in the BI3406-treated group were associated with a partial restoration of β-cell functionality. This observation was confirmed by measuring serum insulin levels in mice from all three groups during IPGTT at D40 (Fig 3D). Here again we demonstrated that total serum insulin levels in the BI-3406-treated group were significantly higher compared to STZ control mice, but did not reach untreated control levels (Fig 3D, E). A similar pattern was observed when measuring serum C-peptide levels, indicating that de novo insulin synthesis was also partially restored in the BI-3406-treated group (Fig 3F). Interestingly, we found that serum glucagon levels remained unchanged between groups (Fig 3G).

To examine the effects of BI-3406 on islet morphology, we harvested pancreata from the three experimental groups at D40 and analyzed them by immunofluorescent staining (Fig 3H, I). We showed that islets from STZ-mice had reduced levels of C-peptide expression, indicating significant loss of β-cells, but that glucagon-expressing α-cells were not affected. As a result, the typical islet architecture involving β-cells clustered at the islet core and surrounded by α-cells was severely disrupted in STZ-treated mice (Fig 3H). Remarkably, BI-3406 treatment led to a significant restoration of c-peptide expression, without however restoring normal islet architecture, further indicating that these were newly-formed β-cells, possibly deriving from surviving β-cells that expanded and surrounded the pre-existing α-cells (Fig 3H). Similarly, expression of mature β-cell markers Nkx6.1 and Pdx1 was severely disrupted in STZ mice but significantly restored upon treatment with BI-3406 (Fig 3H, I). We carefully quantitated each of these markers and observed a 2.24-fold increase in Insulin expression in BI-3406-treated mice compared to STZ-controls (Fig 3J), and a 4.38-fold and 5.42-fold increase in Pdx1 and Nkx6-1, respectively (Fig 3K). In all cases, we observed that the restoration of expression did not reach the levels observed in untreated mice, thus confirming our IPGTT and Insulin ELISA results. We also evaluated the expression of the proliferative marker Ki67, which showed a slight increase, but only in a limited number of islets (Fig 3I, L). This outcome was not surprising since the analysis was conducted at the end of the treatment period, when the expected proliferative effect of BI-3406 had already occurred.

To gain a more qualitative insight of the degree of β-cell restoration upon BI-3406 treatment, we performed FluoClearBABB analysis^41^, a variation of the iDisco technique^42^ that allows examination of large pancreatic fragments using immunofluorescent labelling and 3D image analysis on cleared tissue (Fig 3M; see Methods). We focused our analysis on Nkx6.1 immunolabelling, as a measure of β-cell specification, and confirmed that β-cells are lost in STZ mice, but are largely restored upon BI-3406 treatment (Fig 3M). Importantly, we demonstrate that the typically globular or oval-shaped islet structure is significantly disrupted in STZ mice, and not significantly restored in BI-3406-treated mice (Fig 3M). Regenerated β-cells in this group are not clustered in well-defined structures, as evidenced by an islet volume distribution analysis (probability density function per islet size), demonstrating that treated mice end up with significantly smaller islets compared to untreated controls (Fig S3G). This disruption in islet architecture might partially explain why the newly formed β-cells do not acquire full functionality. In addition to the qualitative analysis, we also quantitated the total volume of Nkx6.1, Glucagon, and Ki67 in pancreata from STZ and STZ+BI-3406 mice and observed a 2.64-fold increase in total Nkx6.1 expression, with no effect on Glucagon and Ki67 expression (Fig S3H). The FluoClearBABB results correlated well with our 2D-signal area quantifications for the same markers on the corresponding samples.

We originally chose to administer BI-3406 before stable hyperglycemia was reached (i.e., at D10), to ensure that a critical mass of β-cells was maintained following STZ treatment. However, we also investigated the role of BI-3406 when administered after the establishment of stable hyperglycemia (i.e., at D17). In this case, only half of the treated mice (50% of n=14) responded to the drug and reverted to normoglycemia (Fig S3I). Nevertheless, this “responder” group demonstrated significantly improved glucose tolerance, in contrast to the “non-responder” group which performed identically to the STZ controls during IPGTT (Fig S3J). We observed a significant correlation between blood glucose levels at the start of treatment and the eventual response outcome; mice that would eventually become responders had an average blood glucose of 264 mg/dl (+/-42.8) at D17, whereas the corresponding levels for eventual non-responders were significantly higher, at 331 mg/dl (+/- 47.8) (Fig S3K). This finding indicates that blood glucose levels may serve as a predictive factor for BI-3406 response outcomes, supporting the idea that a critical β-cell mass is required for the drug to exert its effects.

Finally, we attempted to treat mice with a lower dose of BI-3406 (15mg/kg) both early (D10 onset) and late (D17 onset) or to shorten the treatment length (from 30 to 10 days) using the standard BI-3406 dose (50mg/kg). However, these attempts led to either a partial and non-sustainable decrease in blood glucose levels or showed poor prevalence, only positively affecting a small number of the treated animals (Fig S3L).

### BI-3406 induces β-cell proliferation in diabetic mice

To investigate whether β-cell proliferation sufficiently accounts for the observed β-cell expansion after BI-3406 treatment, we analyzed pancreata from STZ+BI3406 mice treated with EdU and harvested early, as illustrated in Figure 4A. Briefly, BI-3406 was administered twice daily, for a short 7-day period (D10-D16), along with three intraperitoneal doses of EdU (at D10, D12, D14) to label and visualize proliferative cells cumulatively during the initial days of BI-3406 treatment. Blood glucose levels, measured during the treatment period (D1-D16), confirmed that the STZ+BI-3406 group responded to the drug and by D15 already retained lower blood glucose levels compared to diabetic controls, without of course yet reaching the levels of untreated controls (Fig 4B). Consistent with the blood glucose measurements, immunofluorescent analysis at D16 already showed a small (19.1%) but definitive reduction in insulin expression levels in diabetic STZ mice (n=3) compared to untreated controls (n=3), and a clear tendency for restoration of insulin expression in BI-3406-treated mice (n=3) (Fig 4C, S4A). Importantly, the immunofluorescent analysis conducted on D16 revealed an increase in islet EdU^+^ and Ki67^+^ cells in BI-3406-treated animals compared to untreated and STZ-treated controls (Fig 4C-E). Signal quantitation showed an average 8.25-fold increase in the number of proliferative (EdU^+^) islet cells and an average 6.92-fold increase specifically in the number of proliferative β-cells (EdU^+^/Ins^+^) in BI-3406-treated mice compared to STZ-controls (Fig 4D). Similarly, we found an average 7.69-fold increase in the number of Ki67^+^ cells and an average 6.26-fold increase in the number of Ki67^+^/Ins^+^ double-positive cells between the two groups (Fig 4E).

**Figure 4.**
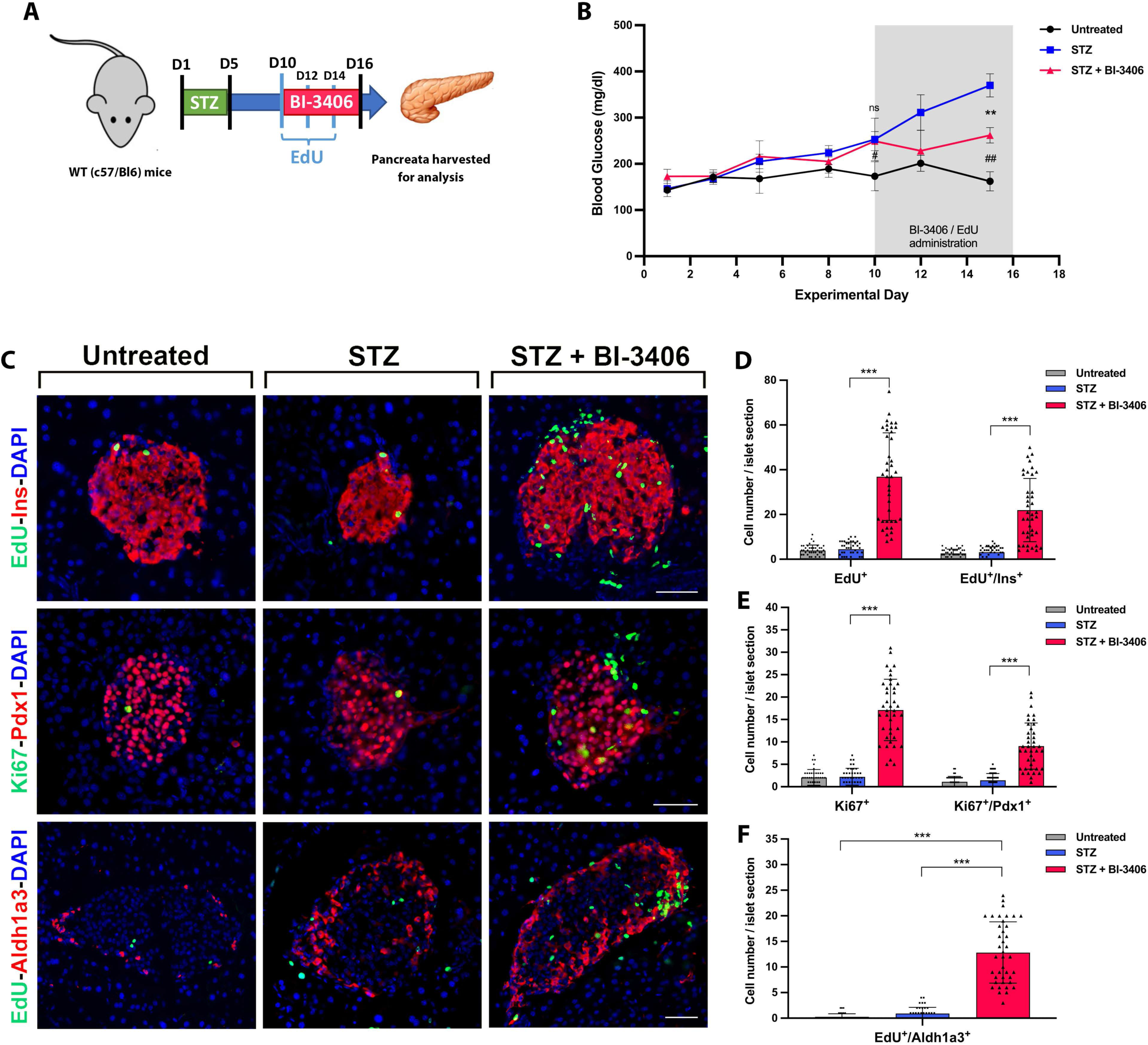
BI-3406 induces β-cell proliferation in diabetic mice. A) Schematic representation of the *in vivo* treatment timeline, including schedule of STZ, EdU and BI-3406 administration. B) Monitoring of blood glucose levels throughout the treatment revealed that, at the time of harvest (D16), mice treated with STZ+BI-3406 (n=3) maintained lower glucose levels compared to those treated with STZ (n=3), although they had not yet reached the normoglycemic levels of untreated mice (n=3). C) Representative images of islet sections from each experimental group, subjected to immunostaining for Insulin, Pdx1, Aldh1a3 and Ki67, as well as fluorescent labelling for the detection of EdU incorporation. D-F) Quantitative assessment of the impact of BI-3406 treatment on islet cell proliferation: Quantitation of the EdU^+^ and EdU^+^/Ins^+^ cells (D), as well as the Ki67^+^ and Ki67^+^/Pdx1^+^ cells (E) revealed a significant expansion in the number of proliferative islet cells, especially β-cells, compared to untreated and diabetic (STZ) controls. F) Quantitation of the number of EdU^+^/Aldh1a3^+^ double-labelled cells indicated a substantial increase in the population of proliferative de-differentiated and immature (Aldh1a3^+^) β-cells, in response to BI-3406. All values are expressed as mean +/-SD. Data points in (B) represent average values per group. Data points in (D-F) represent measurements of individual islet sections per condition (n=40 per group). Data points with zero value are not visible on the x-axis of (D-F). Data were statistically analyzed by one-way ANOVA with Tukey’s post-hoc test. (*) denotes STZ+BI-3406 versus STZ, whereas (^#^) refers to STZ+BI-3406 versus Untreated. \*\**p*<0.01, \*\*\**p*<0.001, ^#^*p*<0.05, ^##^*p*<0.01 and ns (not significant). Scale bars, 40 μm.

It was previously reported by Sachs et al^43^ that STZ-induced diabetes using a “multiple, low dose” STZ scheme similar to ours leads to increased levels of β-cell de-differentiation and functional impairment^43^. De-differentiated β-cells are marked by Aldh1a3 expression and are particularly responsive to pharmacological therapy that triggers their maturation and restores their function^43^. We confirm that our STZ model exhibits increased levels of Aldh1a3 expression (9.1-fold up compared to untreated controls) and that expression remains high, 11.2-fold up compared to untreated controls, following a short treatment with BI-3406 (Fig S4B). In both groups, Aldh1a3+ cells are primarily detected in the periphery of the islets, which is where we detect the majority of the proliferative cells (Fig 4C). Importantly, our quantitations revealed an average 13.9-fold increase in the number of EdU^+^/Aldh1a3^+^ cells in the BI-3406-treated mice compared to STZ controls (Fig 4F), indicating a preferential increase of proliferation in the de-differentiated β-cell population following treatment.

Collectively, our data suggest that in the early stages, BI-3406 treatment triggers proliferation of both mature and immature β-cells, thus adequately explaining the degree of β-cell expansion observed at the end of the full-length (30-day) treatment period.

### BI-3406 induces proliferation of human β-cells transplanted in diabetic mice

We next asked whether BI-3406 can stimulate human β-cell proliferation in vivo, following transplantation of human islets into immunocompromised diabetic mice (Fig 5A). A high dose of STZ (STZ^HI^; D1)^37^ was administered to immunocompromised (NOD-SCID) mice in order to induce acute damage to the majority of the endogenous pancreatic β cells, thereby minimizing their interference with human islets in subsequent treatments. Human islets were then transplanted under the kidney capsule of STZ^HI^ mice (D6; Fig S5A), which were then orally treated with BI-3406 for 30 consecutive days (Fig 5A; details in Methods). We used a small number of human islets (500 IEQ per mouse)^22^, sufficient to maintain stable hyperglycemia but inadequate to fully reverse the diabetic phenotype, in order to allow us to focus on the specific effects of BI-3406.

**Figure 5.**
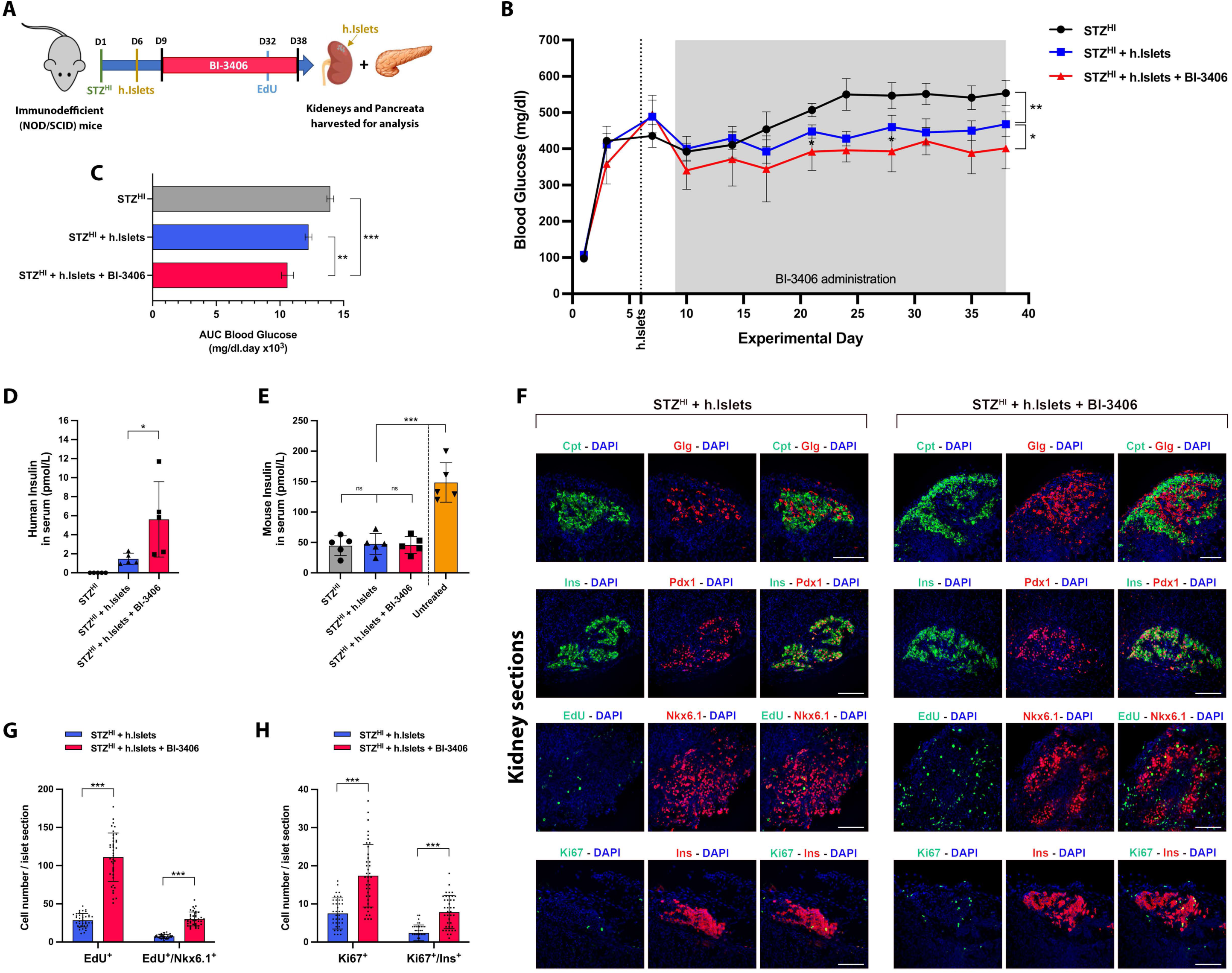
Enhanced β-cell proliferation in human islets transplanted into diabetic mice following treatment with BI-3406. A) Schematic representation of the *in vivo* treatment timeline, including schedule of STZ^HI^, h.Islet, EdU and BI-3406 administration. B) Monitoring of blood glucose levels throughout the treatment demonstrated that mice receiving STZ^HI^+h.Islets+BI-3406 maintained lower glucose levels in comparison to those treated with STZ^HI^+h.Islets or STZ^HI^ alone (n=6 per group). C) Calculations of the areas under the curve (AUC) from the blood glucose monitoring graph further validated that the STZ^HI^+h.Islets+BI-3406 group consistently upheld lower glucose levels throughout the course of BI-3406 treatment, when compared to the control groups. D) An ELISA specific for human insulin, indicated elevated insulin levels in the serum of STZ^HI^+h.Islets+BI-3406 mice, compared to STZ^HI^+h.Islets controls (3.81-fold up), while remaining undetectable in STZ^HI^ mice, as expected. E) Conversely, an ELISA specific for mouse insulin demonstrated a significant decrease in insulin levels in all mice treated with STZ^HI^, with no discernible differences between groups. F) Representative images of kidney sections from STZ^HI^+h.Islets and STZ^HI^+h.Islets+BI-3406 mice, showing transplanted human islets immunostained for C-peptide, Glucagon, Insulin, Pdx1, Nkx6.1 and Ki67, as well as labelled for the detection of EdU incorporation. G-H) Quantitative assessment of the impact of BI-3406 treatment on the proliferation of transplanted human islet cells: Quantitation of the EdU^+^ and EdU^+^/Nkx6.1^+^ cells (G), as well as the Ki67^+^ and Ki67^+^/Ins^+^ cells (H) revealed a significant expansion in the number of proliferative human islet cells, including β-cells, in BI-3406-treated mice compared to controls. Values in (B, D-E, G-H) are expressed as mean +/- SD, and in (C) as mean +/-SEM. Data points in (B) represent average values per group. Data points in (D-E) represent averages from triplicate ELISA measurements per serum sample (n=5 mice per group). Data points in (G-H) represent measurements of individual kidney-embedded islet sections per condition (n=40 per group). Data points with zero value are not visible on the x-axis of (G-H). Data were statistically analysed by one-way ANOVA with Tukey’s post-hoc test (C-E) and by unpaired two-tailed t tests (G, H). \**p*<0.05, \*\**p*<0.01 and \*\*\**p*<0.001. Scale bars, 40 μm.

We monitored blood glucose levels throughout the course of treatment and observed that STZ^HI^ mice (n=6) displayed a rapid surge in blood glucose levels, which continued to rise at a slower rate, finally resulting in stable hyperglycemia in the range of 500-600 mg/dl (Fig 5B). Diabetic mice that received human islets (n=6) displayed an initial decrease in blood glucose levels soon after transplantation and maintained stable hyperglycemia at a lower range (400-500 mg/dl) compared to STZ^HI^ controls (Fig 5B). Importantly, diabetic mice carrying human islets and receiving BI-3406 treatment (n=6) maintained significantly lower blood glucose levels in the range of 350-450 mg/dl compared to controls (Fig 5B, C). This was further supported by evidence that circulating Human insulin in the serum of BI-3406-treated mice showed an average 3.81-fold increase compared to untreated STZ^HI^+h.Islets controls (Fig 5D). We note however that, under these conditions, BI-3406 treatment could not fully revert mice back to normoglycemia, or lead to a substantial improvement in glucose clearance during IPGTT (Fig S5B), likely due to the small number of human islets used for transplantation.

To examine the effects of BI-3406 on endogenous (mouse) and transplanted (human) islets, we harvested pancreata and kidneys from the three experimental groups and performed immunofluorescent analysis. We demonstrated a strong damage in the mouse islets and a remarkable reduction in endogenous β-cell numbers, as expected from the STZ^HI^ dose scheme used, but no significant restoration in response to BI-3406 treatment (Fig S5C, D). Accordingly, we measured the levels of Mouse insulin in the serum of STZ^HI^-treated animals and found a substantial (3.32-fold) reduction compared to untreated controls (Fig 5E). Mouse-insulin levels were unaffected by the presence of human islets and remained equally low even following treatment with BI-3406 (Fig 5E). The fact that BI-3406 treatment did not induce endogenous β-cell expansion supports the argument that a critical β-cell mass is required for the drug to act upon, and also justifies the use of a severely disruptive STZ-dose scheme, in order to prevent endogenous mouse β-cells from competing with the transplanted human β-cells.

In addition to the pancreata, we harvested kidneys from the experimental groups that received islet transplants, and used immunostaining for human endocrine cell-specific markers, to detect the grafted tissue. Human islets in both groups showed strong expression of the α- and β-cell-specific markers glucagon and insulin, respectively, at the expected ratios and expression pattern (Fig 5F). Moreover, we observed strong expression of the mature-β-cell markers Pdx1 and Nkx6-1, indicating that human β-cells maintained their functionality several weeks following transplantation (Fig 5F). To assess proliferation in the transplanted human islets, we used EdU labelling detection and immunofluorescent staining for Ki67 (Fig 5F). We found an average 3.89-fold increase in the total number of EdU+ cells in the human islets of BI-3406-treated kidney samples, including an average 4.06-fold increase specifically in the number of EdU^+^/Nkx6.1^+^ human β-cells (Fig 5G). Interestingly, we observed that the percentage of EdU^+^ cells co-expressing the mature β-cell marker Nkx6.1 is only 27.11% in the BI-3406-treated group, which is significantly lower compared to the *in-vitro* treated islets (51.97%; compare Figs 5G and 2E). Nevertheless, we note that the percentage of Ki67^+^ cells co-expressing Insulin (the generic marker for all types of β-cells) is 45.11% in the BI-3406-treated group (Fig 5H), better reflecting our observations made in cultured human islets. Similar to the EdU measurements, we observed an average 2.33-fold increase in the total number of Ki67^+^ islet cells in response to BI-3406, including an average 3.31-fold increase in the number of Ki67^+^/Ins^+^ cells specifically (Fig 5H).

In conclusion, our findings demonstrate that BI-3406 treatment successfully stimulates proliferation of human β-cells not only in vitro but also in an in-vivo experimental context.

## DISCUSSION

Loss of β-cell mass is a shared characteristic of both type 1 and type 2 diabetes, and various approaches to β-cell replenishment have been explored in mice and humans. Stimulating the proliferation of the remaining pool of endogenous β-cells represents a direct and promising approach for restoring a functional β-cell mass^3–5^.

Multiple mitogenic pathways have been identified as potential targets for β-cell regeneration, however the intentional activation of mitogenic pathways is generally prohibitive in clinical applications due to potential off-target effects that can induce uncontrolled proliferation and tumorigenesis in unrelated tissues. KRAS signaling is generally associated with increased proliferation, however, we establish here that this pathway works in the opposite direction on endocrine cells, and paradoxically inhibits, rather than promotes, β-cell replication. In particular, using appropriate transgenic mouse models, we provide evidence that KRAS overactivation specifically in endocrine progenitors supresses β-cell growth in a cell-autonomous manner, gradually leading to severe hyperglycemia.

With KRAS inhibition being extensively explored as a treatment option for KRAS-driven cancers, we sought to test the hypothesis that the same process may positively impact β-cell proliferation and islet regeneration. Our study demonstrates that the SOS1-RAS inhibitor BI-3406, which is currently under clinical development for various KRAS-driven cancers, indeed fulfils this role.

BI-3406 induces β-cell proliferation in cultured human islets, leading to an increase in the number of EdU- and Ki67-positive cells and overall islet size. The drug is particularly effective in promoting β-cell proliferation, resulting in a significant increase in β-cell mass, while α-cells also respond, albeit to a lesser extent. Crucially, BI-3406 in our hands performed better than Harmine, consistently inducing higher rates of β-cell proliferation across islet samples from multiple donors.

The increase in β-cell mass in human islets was accompanied by improved insulin secretion in response to glucose, suggesting that the newly generated β-cells are functionally more fit, and acquire better glucose sensitivity. Our study revealed that BI-3406 treatment attenuates MAPK signaling, particularly evident by the decrease in MEK1 and ERK1 phosphorylation, while simultaneously displaying a modest but significant increase in STAT3 phosphorylation. STAT3 signaling is associated with anti-apoptotic activity and β-cell protection from DNA damage^44^, suggesting that the induction of β-cell proliferation by BI-3406 may also be linked, directly or indirectly, to β-cell survival.

*In vivo* studies with BI-3406 demonstrated a sustained reversal of hyperglycemia in diabetic mice, associated with an increase in β-cell mass and restoration of insulin secretion. The immediate reduction in blood glucose levels observed in treated mice is most likely attributed to the transient loss of appetite and reduced food intake. However, the surge of β-cell proliferation that occurs in the first week of treatment, followed by the substantial restoration of β-cell mass three weeks later, are primarily responsible for the effective reversion of the diabetic phenotype. Treated mice experience normoglycemia for several days post-treatment, showing improved glucose responsiveness during IPGT tests. However, complete rescue is not achieved, partly because β-cell mass is not fully restored, but also because regenerated β-cells are not reorganized in clusters and therefore do not acquire normal islet architecture which is a prerequisite for full maturation and functionality^45^.

It is evident from our results that BI-3406 requires a pre-existing mass of β-cells to exert its proliferative effect. Mice with severe β-cell loss and blood glucose levels consistently above 250mg/dl do not normally respond to BI-3406, possibly because the surviving pool of responsive β-cells is not adequate. We noticed however, that when a critical islet mass is spared, the injured and partially de-differentiated β-cells in diabetic mice are significantly more responsive to BI-3406 treatment, compared to the healthy and mature β-cells in non-diabetic mice, which show no fluctuations in blood glucose levels during BI-3406 treatment.

Human islets transplanted into immunodeficient diabetic mice also responded positively to SOS1-RAS inhibition, highlighting the potential applicability of this approach as a treatment option for patients with diabetes. Additional pre-clinical studies involving BI-3406 on genetic mouse models of type-2 diabetes (eg the db/db mice), and/or in combination with GLP-1 agonists (eg exenatide, liraglutide etc) to enhance pro-survival activities, would provide additional valuable insights for the most effective clinical use of this approach. Of note, the combination of Harmine with exenatide in pre-clinical testing was shown to enhance the pro-proliferative effects of Harmine by several fold^22^, signifying the necessity to apply a similar combinatorial regimen with the novel SOS1-RAS inhibitor.

As a pan-RAS inhibitor, BI-3406 is anticipated to not only attenuate KRAS signaling but also affect SOS1-HRAS and SOS1-NRAS interactions. While KRAS is considered to be the most important form of RAS operating in β-cells^46^, the potential contribution of HRAS and NRAS inhibition to BI-3406-induced effects cannot be completely ruled out. As new and more potent anti-KRAS-specific drugs emerge, the issue of RAS-specificity will likely be resolved, paving the way for more selective pre-clinical evaluations of these drugs in cancer and other disease indications. For example, two novel drugs developed by Boehringer Ingelheim, BI-2865 and BI-2493, show improved KRAS-specific inhibition activity, with high affinity for the wild-type inactive form of KRAS, while sparing NRAS and HRAS^47^. Our study provides sufficient evidence to warrant the pre-clinical evaluation of these and other similar compounds in various models of diabetes.

In conclusion, understanding the intricate role of KRAS signaling in β-cell proliferation offers significant opportunities for developing novel therapeutic strategies to combat diabetes. The insights gained from this study put into prominence the option of using KRAS as a therapeutic target, and pave the way for new and innovative therapies that will promote β-cell regeneration.

## METHODS

### Mouse strains, maintenance and genotyping

Animal maintenance and experimentation was performed in accordance with the FELASA guidelines for the care and use of laboratory animals (Article 23 of the Directive 2010/63/EU). All procedures conducted were in compliance with the institutional guidelines and approved by the Directorate of Agriculture and Veterinary Policy, Region of Attika, Greece. Mice were housed on a 12hr light/dark cycle in controlled-climate rooms (21-23°C) and were screened on a regular basis to confirm a pathogen-free environment.

Mouse mutant and transgenic lines NOD.Cg-Prkdc^scid^/J (#001303), Tg^(Neurog3-cre)C1Able/J^ (#005667), Tg^(Pdx1-cre)6Tuv/J^ (#014647) and Gt(ROSA)26Sor^tm4(ACTB-tdTomato,-EGFP)Luo/J^, (#007576), were obtained from The Jackson Laboratory (Bar Harbor, Maine). The Actb^tm1(KrasG12D)Arge^ mouse line was a gift from A. Efstratiadis^38^. All experiments involving wild-type mice were performed on C57BL/6J mice. All mouse strains used were interbred onto the C57BL/6J genetic background, with the exception of NOD.Cg-Prkdc^scid^/J (NOD/SCID) mice. Genotyping was performed by conventional PCR on genomic DNA isolated from mouse-tails using standard procedures. Briefly, mouse tails were dissolved in Tail buffer (100 mM Tris-HCl pH 8.0, 200 mM NaCl, 5mM EDTA, and 0.2% SDS) supplemented with 50 μg/ml Proteinase K (Sigma) for 16 hrs at 55 C°. Following protein extraction with a 1:1 Phenol/Chloroform mixture (Sigma), genomic DNA was precipitated from the aqueous phase with 100% ethanol and finally resuspended in TE buffer (10 mM Tris-HCl pH 8.0 and 1 mM EDTA). PCR primers for genotyping are provided in Table S1.

### STZ treatments

For the “multiple; low-dose” STZ scheme, 8-week-old male C57BL/6 mice were injected intraperitoneally with 40mg/kg streptozotocin (STZ) (Sigma S0130) dissolved in 50mM citrate buffer (50mM Na-citrate, 50mM citric acid, pH 4.5), once daily for 5 consecutive days. A subset of age-matched control mice was injected with plain citrate buffer. Throughout the course of treatment, mice were fasted for 4 hours prior to STZ injection and were provided with water supplemented with 10% sucrose, which was replaced with regular water 24 hours after the last injection. Blood glucose levels were regularly measured as described below, to ensure that stable hyperglycemia was reached, normally 8-10 days after the last STZ injection. For the “single; high-dose” STZ scheme (STZ^HI^), 8-week-old male NOD/SCID mice were fasted for 4 hours and subsequently received a single intraperitoneal injection of 180mg/kg STZ, dissolved in 50mM citrate buffer (pH 4.5). Treated mice were provided with 10% sucrose water and closely monitored for 12 hours, for marked hypoactivity or other adverse reactions. Blood glucose levels were regularly measured thereafter to ensure that stable hyperglycemia was reached, normally 48 hours after STZ administration.

### Formulation and administration of BI-3406

The vehicle used for the *in vivo* delivery of BI-3406 was 0,5% Natrosol 250Hx Hydroxyethylcellulose (Ashland), dissolved in water and prepared as described in ^35^. Briefly, 200ml water was gradually added to 1g of Natrosol and subsequently heated at 50°C for 5 hours. The solution was left stirring overnight at RT and subsequently autoclaved for 20 min at 121-125 °C, 100 bar. BI-3406 solution was formulated by slowly adding 0,5% Natrosol to the weighed compounds at the desired doses, after correcting for the indicated purity factors. Solutions were stirred at 700-1000 rpm for 3 hours at RT, then sonicated for 3 min at 35kHz in an ultrasound bath, and finally the pH adjusted to 4.5 with 1M HCl. Immediately prior to delivery, the formulated compounds were subjected to a final sonication step to ensure a homogeneous suspension.

Wild-type and STZ-treated mice were randomized in experimental and control groups. Experimental groups received BI-3406 at 15 or 50mg/kg as indicated, whereas control groups received the vehicle (0.5% Natrosol) alone. Drugs and vehicle were administered by oral gavage, twice per day with a 6-hour interval, for 30 consecutive days, unless otherwise stated.

### Blood glucose monitoring and hormone detection

Blood glucose levels and body weight were regularly monitored (2-3 times per week) in all animals of every experimental and control group, throughout each course of treatment. Following a 4-hour fasting, animals were weighed and glucose concentration was measured from a tail-vein blood drop using a handheld glucometer (Contour Next; Bayer) according to the manufacturer’s instructions.

For hormone measurements in serum, whole blood was collected from the submandibular vein of 4-hour fasted animals, allowed to clot by standing for 30 minutes at room temperature and centrifuged at 13000rpm for 12 minutes to pellet cells. Insulin (Mercodia; 10-1249), C-peptide (Chrystal Chem; 90050) and Glucagon (Mercodia; 10-1281) levels were assayed in the supernatant using appropriate ELISA kits and following the manufacturer’s instructions.

### Intraperritoneal glucose tolerance test (IPGTT)

Mice were fasted for 16h overnight and subsequently injected intraperitoneally with D-glucose (20% w/v solution; Sigma) at 2 g/kg. Blood from tail vein was used to determined glucose levels at t0 and at 15, 30, 60, 90 and 120 min following glucose injection, using a high-range handheld glucometer (AlphaTRAK 2; Zoetis). For insulin-during-IPGTT measurements, tail vein blood samples were collected at the indicated time-points using heparinized capillary tubes (Fisher), allowed to clot by standing for 30 minutes at room temperature and centrifuged at 13000 rpm for 12 minutes to pellet cells. Insulin was assayed in the supernatant using an ultrasensitive mouse insulin ELISA kit (Mercodia).

### EdU administration and detection

The modified uracil analogue 5’ethynyl-2’-desoxyuridine (EdU) was used to investigate cell proliferation. EdU (Sigma) was injected intraperitoneally at 30 mg/kg body weight at the indicated time-points prior to sacrifice. EdU incorporation in cells was detected by immunofluorescence on tissue cryosections using the Click-It EdU Cell Proliferation kit (Invitrogen), following the manufacturer’s instructions.

### Culture and pharmacological treatment of human islets

Primary human cadaveric islets were obtained from Prodo Labs Inc. (Irvine, CA, USA). Prodo Labs offers live Human Islets for Research (HIR), isolated from non-diabetic cadaveric donors that perished from head trauma or other non-pathological cause, and accompanied by formal written informed consent. In total, islets from 6 donors were obtained (3-10,000 IEQ per donor depending on availability), spanning a wide range of ages and BMIs as depicted in Fig 2B. Upon receipt, islets were transferred to non-adherent P10 tissue culture plates and allowed to recover in complete PIM(R) medium [PIM(R) supplemented with 5% Human Serum PIM(ABS), 2mM Glutamine PIM(G) and antibiotics mix PIM(3X)] (all from Prodo Labs), for 48h at 37C and 5% CO_2_. Islets where then transferred in 24-well suspension plates, in complete PIM(S) medium (Prodo Labs) at 100 IEQ per well, and were treated with BI-3406 (100nM-5μΜ) or Harmine (10μΜ; Sigma), both dissolved in DMSO (<0.1% v/v). Control islets were treated with plain DMSO. Islets were cultured for 7 days, and drugs were replenished every second day with each medium change. EdU (Invitrogen) was added at 10 μg/ml with the last medium change, 48h before islet harvesting. We initially tested a range of BI-3406 concentrations on Donor-1 islet cell proliferation, and established that 300nM works optimally (Fig S2A), therefore this concentration was used in all subsequent *in vitro* assays. Nevertheless, we found that BI-3406 exerts no adverse effects on islet cell viability, even at 5μM, the highest concentration tested (Fig S2B).

### Glucose-stimulated insulin secretion (GSIS)

Following treatment, human islets were washed in ECS (125mM NaCl, 2.5mM KCl, 26mM NaHCO_3_, 1.25mM NaH_2_PO_4_, 1mM MgCl_2_, 2mM CaCl_2_ and 10mM Hepes pH 7.4), and transferred by hand-picking in a round-bottom 96-well plate at 10 IEQ/well (at least 3 wells per donor) in 200μl ECS supplemented with 3mM D-glucose (Sigma). Islets were equilibrated for 1 hour at 37°C and 5% CO_2_ and medium was replaced with fresh ECS containing 3mM D-glucose (baseline medium). Islets were incubated for 1h at 37°C and 5% CO_2_, and baseline medium was collected and replaced with stimulatory medium (ECS with 20mM D-glucose) for another hour at 37°C and 5% CO_2_. Insulin secreted in the baseline and stimulatory media was measured using a human Insulin ELISA (Chrystal Chem), following the manufacturer’s instructions. Results were normalized for total protein, extracted by *in situ* lysis of islets in RIPA buffer (50 mM Tris-HCl pH 7.4, 1% NP-40, 0.5% Na-deoxycholate, 0.1% SDS, 150 mM NaCl, and 2 mM EDTA), supplemented with protease and phosphatase inhibitor cocktails (Sigma), and quantitated using the QuantiPro BCA Assay kit (Sigma).

### Transplantation of human islets

Human islets from Donor 6 were transplanted into diabetic NOD/SCID recipient mice. Recipients were made diabetic following the “single; high-dose” STZ scheme, and islet transplantation was performed 5 days after the single STZ injection, as described in ^48^. Briefly, recipient mice were anesthetized with 3% sevoflurane in a 4% O_2_ chamber, shaved and cleaned with betadine/alcohol solution. A 1cm incision was made at the ventral site of the body, and the kidney was exposed after applying light pressure. Human islets (500 IEQ resuspended in 10μl PBS) were gently injected under the kidney capsule (Fig S5A), using a custom-made catheter needle. The kidney was placed back into the abdominal cavity, the peritoneum was sewed with 4-0 absorbable continuous suture (Vicryl Suture; Ethicon) and the skin was stapled with surgical clips (9mm AutoClip; FST). Mice were left to recover on a heating blanket and Meloxicam (0.2mg/kg; Dopharma) was applied for pain relief. Mice were closely monitored for post-operative adverse reactions, and BI-3406 treatment was initiated 3 days after islet transplantation (Fig 5A).

### Immunostainings on cryosections and histological analysis

Pancreata and kidneys from euthanized mice, and cultured human islets, were collected, fixed in 4% paraformaldehyde (PFA) (Sigma) at 4^0^C for 30 min (human islets) or 2h (mouse tissues), washed in PBS and dehydrated in 30% sucrose (sucrose) o/n at 4^0^C. Samples were embedded in optimal cutting temperature (OCT; Sakura), cut into 12μm-thick sections and mounted onto superfrost slides (Thermo Scientific) for storage at -80C. Embryonic (E14.5 and E16.5) pancreata were harvested and processed as described in ^49^. For immunostainings, cryosections were washed with PBS and blocked for 1 hour at RT in 10 x blocking solution (10% normal goat serum and 0.3% Triton X-100 in PBS). Primary antibodies were diluted in 1 x blocking solution and incubated o/n at 4 °C, whereas secondary antibodies, also diluted in 1 x blocking solution, were incubated for 1h at room temperature. All washes were done in PBST (PBS with 0.3% Triton X-100). After immunostaining, slides were mounted with DAPI (Vectashield; Vector) and taken for fluorescent microscopy. A list of primary and secondary antibodies used is provided in Table S2.

For histological analysis, mouse pancreata were dissected, fixed in 10% formalin (Sigma) followed by dehydration in 70% ethanol, embedded in paraffin, cut into 6 μm thick sections and mounted onto polylysine-coated slides (Thermo Scientific). Sections were then stained with hematoxylin (30s at RT) and eosin (60s at RT), according to standard procedures. Slides were left to dry and covered with DPX mounting medium (Sigma) for examination under brightfield light microscopy.

### Morphometric analysis of immunofluorescent images

Morphometric analysis and signal quantitation on adult mouse tissues (pancreata and kidneys) and human islets was performed as described^49^, using immunofluorescent images taken at constant settings, and the Image J (Fiji) software. More specifically, for the endocrine markers: Insulin, Glucagon and Aldh1a3, the “Analyse Particles” tool with standard thresholds across samples was used to analyse the number of pixels that correspond to signal area on an immunofluorescent image, whereas for the nuclear markers: EdU, Ki67, Pdx1 and Nkx6.1, signal-positive nuclei were counted directly on the immunofluorescent image using the “Cell Counter” plugin. For signal normalisations over DAPI, this was either measured as total signal area in pixels (for entire tissue sections), or by nuclei counting (when individual islets were analysed).

For transgenic mouse lines AKras* and Ngn3Cre-AKRas*, Insulin and Glucagon signal areas were expressed as a percentage of total DAPI signal area per field, and analysis was performed using at least twelve 12-μm-thick cryosections per genotype, collected 100μm apart to span the entire tissue and ensure that any given islet or endocrine cell cluster is scored only once. For drug-treated and control animals, Insulin, Glucagon and Aldh1a3 expression was presented as signal area per islet section, whereas EdU, Ki67, Pdx1 and Nkx6.1 expression was presented as the number of signal-positive nuclei per islet section. Analysis was performed on 12-μm-thick pancreas or kidney cryosections, collected 100μm apart to span the entire tissue, aiming to score at least 40 islet structures per condition, after excluding the strong outliers, defined as those with signal areas higher than the 3^rd^ quartile plus the tripled interquartile range (3.0xIQR) or lower than the 1st quartile minus 3.0xIQR. For all quantitations involving mouse tissue sections, at least three tissues (pancreata and/or kidneys) of each genotype or condition were used.

For human islets, the number of EdU^+^ or EdU^+^/Nkx6.1^+^ nuclei were counted and expressed as a percentage of total DAPI^+^ or Nkx6.1^+^ nuclei per islet section. A total of 40-60 islets were analysed per condition and per donor, after excluding the strong outliers identified as described above. Islet mass was expressed as the average number of DAPI^+^ nuclei per islet section for each donor, whereas β-cell mass was expressed as the average number of Nkx6.1^+^ nuclei per islet section for each donor.

### FluoClearBABB and image analysis

FluoClearBABB, a variation of iDisco, was used for the fluorescent immunolabeling of large pancreatic tissue samples, for volume imaging and analysis^41,42^. Following perfusion, whole pancreatic tail fragments from BI3406-treated and control mice (3 per group) were dissected and fixed in 4% PFA in PBS for 24h at 4C. Staining of the pancreatic fragments was performed as previously described^42^, using the following fluorophore-conjugated primary antibodies: Nkx6.1 (Cell Signaling) labelled using the AlexaFluor-700 conjugation kit (Abcam), Glucagon (Sigma) labelled with AlexaFluor-488 (Abcam) and Ki67 (Abcam) labelled with AlexaFluor-405 (Abcam). Stained tissue fragments were cleared using the FluoClearBABB protocol^41^ and images were acquired using a Leica Stellaris 8 microscope equipped with a 20x BABB immersion lens (HCX APO L 20×/0.95 IMM) and a white light laser. For image visualisation, a non-signal channel was acquired and used to render the volume of the pancreatic fragment, using the surface function in Imaris (Bitplane; surface grain size 14 µm; manual threshold of 1 and volume above 3,97e7 µm3). To show the distribution of Nkx6.1^+^ islets within the tissue, the Nkx6.1 signal was rendered using the surface function and a surface grain size of 2.3µm, manual threshold of 34.59 and filter “volume” above 166 µm^3^.

Signal volumes were computed using the surface function in Imaris (Bitplane) with the following parameters: Nkx6.1 (surface grain size of 2.1µm; manual threshold value of 25.57; filter “volume” above 166µm^3^), Glucagon (surface grain size of 2.1µm; manual threshold value of 6.52; filter “volume” above 166µm^3^), Ki67 (surface grain size of 2.1µm; manual threshold value of 1.14; filter “volume” above 166µm^3^), Total volume (surface grain size of 15µm; manual threshold value of 1; filter “volume” above 3.97e7µm^3^). For islet size distribution analysis, islet volumes were calculated using Nkx6.1 immunofluorescence, and distributions were expressed as the probability density function, following the procedures described in^50^. Signal volume calculations, including average values, standard deviations and statistical analyses, were performed using MS Excel, based on the “volume” attribute supplied by Imaris.

### Phosphoprotein multiplex immunoassay

DMSO- and BI3406-treated human islets from a single donor (Donor 6) were used for the relative quantitation of the phosphorylation levels of 20 phosphoproteins involved in major signaling pathways, using the Luminex FlexMap3D platform. Briefly, treated and untreated human islets (1500 IEQ per sample) from Donor 6, were lysed following Protavio’s Lysis protocol, and protein concentration was determined using the QuantiPro BCA Assay kit (Sigma) and adjusted at 500 μg/ml. Assays were performed by Protavio Ltd, which specializes on the xMAP technology developed by Luminex. Using advanced fluidics, optics and digital signal processing, this technology allows the simultaneous measurement of multiple biochemical markers in a very small sample volume. Assays were therefore performed in 35μm sample volumes, multiplexed into analyte-specific color-coded micropsheres to attain distinct fluorescent signatures, as described in. Protavio’s in-house custom phosphoprotein assay panel #HSIGNAL-20 was used for sample testing, containing the following biomarkers: SMAD3, P53, AKT1S1, GSK3, AKT1, HSPB1, P38, MEK1, RSK1, CREB1, IKBA, MTOR, JUN, EGFR, ERK1, PTN11, STAT3, CHK2, NFKB and MARCKS. Samples were run in triplicates and lysates from different cell lines were used as positive and negative controls. Signal to noise ratio (SNR) was calculated for each assay, and final results were presented as average Median Fluorescence Intensity (MFI) from the three replicate measurements for each analyte in each sample.

### Western blotting

Protein extraction, gel electrophoresis, and immunoblotting were conducted following standard protocols. For protein extraction, samples were lysed in RIPA buffer (Millipore), supplemented with protease and phosphatase inhibitor cocktails (Sigma). Proteins were quantitated using the QuantiPro BCA Assay kit (Sigma) and loaded at 30 μg/lane on a polyacrylamide gel for electrophoresis. Protein transfer was performed on a standard nitrocellulose membrane (Amersham), which was subsequently stained with Ponceau S (Sigma). Blocking was performed in 5% milk in TBST (50 mM Tris-HCl pH 7.6, 150 mM NaCl, and 0.05% Tween-20) for 1 h at room temperature, and primary and secondary antibodies were diluted in 1% milk in TBST and incubated overnight at 4 C° or for 2 h at room temperature, respectively. Signal was detected with chemiluminescence (ECL; Thermo Fisher) and developed on an Optimax 2010 X-Ray Film Processor. Primary antibodies used were mouse anti-B-Raf (1:1000; Cell Signaling), rabbit anti-Phospho-B-Raf (1:1000; Cell Signaling) and mouse anti β-actin (1:5,000; Santa Cruz). Secondary antibodies were anti-rabbit and anti-mouse horseradish peroxidase (HRP)-conjugated goat antibodies (1:5,000; Dako).

### Statistical analysis

Data are expressed as means and presented in line charts, box plots or bar graphs generated using the GraphPad Prism Software (v. 9). Error bars represent Standard Deviation (SD) unless otherwise stated. Statistical significance was determined by Student’s t tests for two-tailed distribution of unpaired groups, or by one-way ANOVA followed by Tukey’s post hoc tests, and differences with p<0.05 were considered significant. For differences in islet size distributions (iDisco analysis), statistical significance was determined using the Wilcoxon rank-sum test. The sample size (n) for each experiment is indicated in the corresponding figure legend and, unless otherwise stated, it refers to biological replicates.

## ACKNOWLDEGMENTS

This work was financially supported by the “OpnMe” Innovation Portal action “Molecules for Collaboration” (CRA No: 493341 to I.S.), awarded by and conducted in collaboration with Boehringer Ingelheim’s Research Beyond Borders (RBB) team. We thank Argiris Efstratiadis (BRFAA) for his generous gift of the Actb^LSLKras*^ mouse line and for critical reading of the final version of the manuscript. We thank Pavlos Alexakos and Vagelis Balafas at BRFAA’s animal house for expert animal husbandry and assistance with critical mouse handling procedures. Finally, we are grateful to the donors and their families who permitted the use of human islets, provided to us via Prodo Labs.

## AUTHOR CONTRIBUTIONS

I.S., G.W. and T.K. conceived the study and designed the experiments. A.P., C.C., P.K. and D.R. performed the investigation. A.P., J.S., A.G., M.H., M.F. J.D., P.E., T.K., G.W. and I.S. conducted the analysis and interpretation of results. I.S. wrote the original draft, with critical revision from A.G., G.W., T.K. and M.H. All authors reviewed, edited and approved the final manuscript. I.S. takes primary responsibility for the data described in this manuscript.

