## Supplement for "Pharmacological targeting of SOS1-RAS interaction triggers pancreatic β-cell proliferation and sustainably reverses diabetic hyperglycemia"

#### SUPPLEMENTARY FIGURE LEGENDS

**Figure S1**

KRAS overactivation in the pancreatic epithelium and validation of the Ngn3Cre mouse line. A) Pdx1Cre-AKras* mice that express a mutated (KRAS$^{G12D}$) and constitutively active form of KRAS, under the control of the β-actin promoter, exclusively in the pancreatic epithelium (Pdx1$^+$ cells), were analysed and compared to unrecombined AKras* controls. B) Fasting blood glucose levels at 4 weeks were significantly increased in Pdx1Cre-AKras* mice (n=8) compared to age-matched controls (n=8). C) Ngn3Cre-RosaGFP mice were used to permanently label Ngn3$^+$ cells and their progeny with mEGFP, in order to verify the specificity and fidelity of the Ngn3Cre transgenic line. D) Representative images of mEGFP-Ngn3 (E14.5) and mEGFP-E.Cadherin (E16.5) co-expression indicate that Ngn3+ cells and their progeny are specifically labelled in the Ngn3Cre-RosaGFP line during embryonic development, leading to adult pancreata with strong mEGFP fluorescence exclusively in the endocrine compartment (n=3 for all groups). Values in (B) are expressed as mean +/-SD, and data were analysed by an unpaired two-tailed t test (***$p$<0.001). Scale bars, 80 µm.

**Figure S2**

Evaluating the impact of BI-3406 on cultured human islets. A) Dose-response analysis of BI-3406 on β-cell proliferation in human islets. Quantitation of the number of EdU$^+$/Nkx6.1$^+$ cells (expressed as a percentage of total EdU+ cells) accross a range of BI-3406 concentrations (0, 0.1, 0.3, 1 and 5 µM) suggested an optimal concentration of 300 nM, which was subsequently utilized in all in vitro experiments. B) No detrimental effects were observed in response to BI-3406 treatment, even at the highest concentration tested (5 µM), as evidenced by the absence of necrotic cells (normal nuclei morphology after DAPI staining), robust Insulin expression and the absence of pro-apoptotic (Casp3$^+$) cells. Extended cultures of DMSO-treated islets (>3 weeks) were used as a positive control for deteriorating islets, characterised by pronounced Casp3 expression at the islet core and a marked reduction in Insulin expression. C) Representative images of islet sections from Donor 6 following treatment with BI-3406 or DMSO, immunostained for Glucagon (Glg) and labelled for the detection of EdU incorporation. D) Quantitation of the number of EdU$^+$/Glg$^+$ cells (expressed as a percentage of total EdU$^+$ cells) unveiled a modest yet significant increase (1.99-fold) in α-cell proliferation upon BI-3406 treatment. E) Plot illustrating the fold-increase in proliferative (EdU$^+$) cells in response to BI-3406 for each donor, relative to their body-mass index (BMI), revealing a linear correlation with an $R^2$ value of 0.8602. F) Comparison of the effects of BI-3406 and Harmine on β-cell proliferation (EdU$^+$/Nkx6.1$^+$ cells expressed as a percentage of total Nkx6.1+ cells), evaluated under identical conditions, demonstrated superior performance by BI-3406 for every donor tested accross a broad age range (24.8% higher β-cell proliferation for Donor 1, 34.4% for Donor 3 and 31.1% for Donor 6). G) Plot of absolute fluorescence intensity values (Luminex assay) corresponding to the expression profiles of 20 key phosphoproteins detected in the lysates of DMSO- and BI-3406-treated islets. All values are expressed as mean +/-SD. Data points in (A) represent average values of individual islet measurements (n=40 per BI-3406 concentration). Data points in (D) represent measurements of individual islets from Donor 6 (n=40 per condition). Data points with zero value are not visible on the x-axis of (D). Data points in (E) represent average values per donor. Data were statistically analysed by unpaired two-tailed t tests (A, D, G) and one-way ANOVA with Tukey's post-hoc test (F). $^{*}p<0.05$, $^{**}p<0.01$ and $^{***}p<0.001$. Scale bars, 40 μm.

**Figure S3**

Monitoring the effects of BI-3406 treatment in vivo. A) Monitoring of mouse body weight throughout the treatment period, revealed a modest weight reduction in STZ+BI-3406 mice upon initiation of BI-3406 administration, which was completely reversed by the end of treatment. B-D) Monitoring of blood glucose levels for untreated (B), STZ (C) and STZ+BI-3406 (D) mice, demonstrating fluctuations throughout treatment for each individual animal. E) Wild-type animals treated with standard dose BI-3406 (50mg/kg), showed no significant effect on blood glucose levels compared to untreated animals. F) IPGT tests conducted on untreated, STZ and STZ+BI-3406 mice at the end of the post-treatment period (D60), revealed that blood glucose clearance remained significantly more efficient in BI-3406-treated mice compared to STZ controls, however, these rates still did not reach the levels observed in untreated controls (n=5 per group). G) Probability density function plot of islet volume, calculated from volume data derived from FluoClearBABB (iDisco) image analysis, demonstrated that the median volume of the regenerated islets in the STZ+BI-3406 group ($20.2 \times 10^3$ μm$^3$) is significantly smaller compared to that of the untreated controls ($41.1 \times 10^3$ μm$^3$). H) FluoClearBABB image analysis showed a significant restoration of Nkx6.1 signal volume in BI-3406-treated pancreata compared to STZ controls, with no observable effect on Glucagon and Ki67 signal volume. I) Late initiation of BI-3406 treatment (D17) resulted in a mixed group consisting of responders (n=7) who revert to normoglycmia, and non-responders (n=7) who retained high blood glucose levels throughout treatment. J) IPGT tests at the end of the late-initiated BI-3406 treatment, revealed a clear distinction between responder and non-responder groups, with the former demonstrating significantly improved glucose tolerance (n=5 per group). K) Comparisons of blood glucose level between responders and non-responder groups at various stages of treatment revealed that lower glucose levels at treatment initiation correlated with a positive response at the end of treatment (n=7 per group). L) Monitoring of blood glucose level fluctuations in mice treated with a lower dose of BI-3406

(15mg/kg) either early (D10 onset) (n=6; Responders=4, Non-responders=2) or late (D17 onset) (n=6), or with the standard dose (50mg/kg) over a shorter period (10 days instead of 30) (n=6; Responders=3, Non-responders=3), revealed a mixed response outcome. All values are expressed as mean +/-SD. Data points in (A, E-F, H-L) represent average values per group. Data points in (B-D) represent glucose measurements of individual mice per time-point. Data points in (H) represent average signal volume measurements over total tissue volume per condition, expressed as a percentage of untreated (n=3 per group). Data were statistically analysed by one-way ANOVA with Tukey's post-hoc test (A, E-F, I-J, L), by unpaired two-tailed t tests (H, K) and by Wilcoxon rank-sum test (G). (*) denotes STZ+BI-3406 versus STZ, whereas ($^{\#}$) refers to STZ+BI-3406 versus Untreated (F, J). **p*<0.05, ***p*<0.01, ****p*<0.001, $^{\#}p<0.05$, $^{\#\#}p<0.01$, $^{\#\#\#}p<0.001$ and ns (not significant).

**Figure S4**

Short-term BI-3406 treatment does not affect islet morphology. A) Quantitative analysis of the signal area corresponding to Insulin expression confirmed a reduction in the number of β-cells, however, a brief 7-day treatment with BI-3406 did not result in a significant restoration of β-cell numbers. B) Quantitative analysis of the signal area corresponding to Aldh1a3 expression demonstrated that STZ-induced diabetes is associated with an increase in the population of de-differentiated and immature β-cells. Notably, this alteration remains unaffected by the short-term treatment with BI-3406. Representative images from the Insulin and Aldh1a3 immunostainings used to generate these quantitative data, can be found in Fig 4C. All values are expressed as mean +/-SD. Data points represent measurements of individual islet sections (n=40 per condition), derived from 3 mice per group. Data were statistically analyzed by one-way ANOVA with Tukey's post-hoc test. **p*<0.01, ***p*<0.001, and ns (not significant).

**Figure S5**

Effect of BI-3406 treatment on endogenous islets in $STZ^{HI}$ diabetic mice transplanted with human islets. A) Example of a mouse kidney that hosts human islets, injected at 500 IEQ (Islet Equivalent: refers to) under the renal capsule, at the site indicated by the black arrow. B) Presence of human islets in $STZ^{HI}$ mice results in a significant improvement in blood glucose clearance during IPGTT, however, BI-3406 treatment does not confer any additional improvement (n=3 per group). C) Representative images of endogenous pancreatic islet sections from each experimental group (n=3 per group), immunostained for Insulin and Glucagon, demonstrate a profound disruption in the islet architecture and morphology across samples. D) Quantitative analysis of the number of $Ins^{+}$ and $Glg^{+}$ cells in these pancreatic samples confirms the substantial reduction in β-cell numbers (though not α-cells) in $STZ^{HI}$ mice, which is not restored by BI-3406 treatment. All values are expressed as mean +/-SD.

Data points in (B) represent average values per group. Data points in (D) represent measurements of individual islet sections per condition (n=20 per group), expressed as a percentage of untreated controls. Data were statistically analyzed by one-way ANOVA with Tukey's post-hoc test. (*) denotes $STZ^{HI}$+h.Islets versus $STZ^{HI}$, whereas ($^{\#}$) refers to $STZ^{HI}$+h.Islets+BI-3406 versus $STZ^{HI}$. *$p<0.05$, **$p<0.01$, ***$p<0.001$, $^{\#}p<0.05$, $^{\#\#}p<0.01$ and ns (not significant). Scale bars, 40 μm.

#### Supplementary Table 1: Genotyping Primers

| | |
|---|---|
| **Neurog3-Cre** | |
| Fw | AAAATTTGCCTGCATTACCG |
| Rv | ATGTTTAGCTGGCCCAAATG |
| **Pdx1-Cre** | |
| Fw | CTGCCACGACCAAGTGACAGC |
| Rv | CTTCTCTACACCTGCGGTGCT |
| **Gt(ROSA)26Sortm4(ACTB-tdTomato,-EGFP** | |
| Fw | CCTACGGCGTGCAGTGCTTCAGC |
| Rv | CGGCGAGCTGCACGCTGCCGTCCTC |
| **Kras$^{G12D}$** | |
| Fw | TGTATGGGATCTGATCTGGGGCCTCG |
| Rv | CTCGGCAGGAGCAAGGTGAGATG |

**Supplementary Table 2: Primary and secondary antibodies**

| Antigen | Clone | Source | Cat. No. | Dilution |
|---|---|---|---|---|
| Aldh1a3 | Rabbit polyclonal | Novus | NBP2-15339 | 1:300 (IF) |
| β-actin | Mouse monoclonal, AC-15 | Sigma-Aldrich | A5441 | 1:10000 (WB) |
| BRAF | Mouse monoclonal, E3T5C | Cell Signaling | 77622 | 1:1000 (WB) |
| Caspace3 (cleaved) | Rabbit polyclonal | Cell Signaling | 9661 | 1:400 (IF) |
| c-peptide | Rabbit polyclonal | Cell Signaling | 4593 | 1:150 (IF) |
| E-cadherin | Rat monoclonal, ECCD-2 | Invitrogen | 13-1900 | 1:400 (IF) |
| Glucagon | Mouse monoclonal, K79bB10 | Sigma-Aldrich | G2654 | 1:500 (IF) |
| Insulin | Mouse monoclonal, K36AC10 | Sigma-Aldrich | I2018 | 1:500 (IF) |
| Ki67 | Rabbit polyclonal | Abcam | ab 15580 | 1:300 (IF) |
| Neurogenin3 | Rabbit polyclonal | Gift from H. Edlund | | 1:200 (IF) |
| Nkx6.1 | Rabbit polyclonal | Cell Signaling | 54551 | 1:400 (IF) |
| Pdx1 | Guinea pig polyclonal | Abcam | Ab 47308 | 1:150 (IF) |
| Pdx1 | Rabbit polyclonal | Gift from C. Wright | | 1:5000 (IF) |
| Phospho-BRAF | Rabbit polyclonal | Cell Signaling | 2696 | 1:1000 (WB) |
| Anti-Mouse IgG AlexaFluor-568 | Goat polyclonal | Invitrogen | A-11019 | 1:500 (IF) |
| Anti-mouse IgG AlexaFluor-488 | Goat polyclonal | Invitrogen | A-11017 | 1:500 (IF) |
| Anti-Rabbit IgG AlexaFluor-568 | Goat polyclonal | Invitrogen | A-11011 | 1:500 (IF) |
| Anti-Rabbit IgG AlexaFluor-488 | Goat polyclonal | Invitrogen | A-11070 | 1:500 (IF) |
| Anti-rabbit IgG AlexaFluor-647 | Goat polyclonal | Invitrogen | A-21244 | 1:500 (IF) |
| Anti-rat IgG AlexaFluor-647 | Goat polyclonal | Invitrogen | A-21247 | 1:500 (IF) |
| Anti-guinea pig IgG AlexaFluor-568 | Goat polyclonal | Invitrogen | A-11075 | 1:500 (IF) |
| Anti-mouse IgG HRP | Horse polyclonal | Cell Signaling | 7076 | 1:5000 (WB) |
| Anti-Rabbit IgG HRP | Goat polyclonal | Cell Signaling | 7074 | 1:5000 (WB) |

Supplemental Figure 1

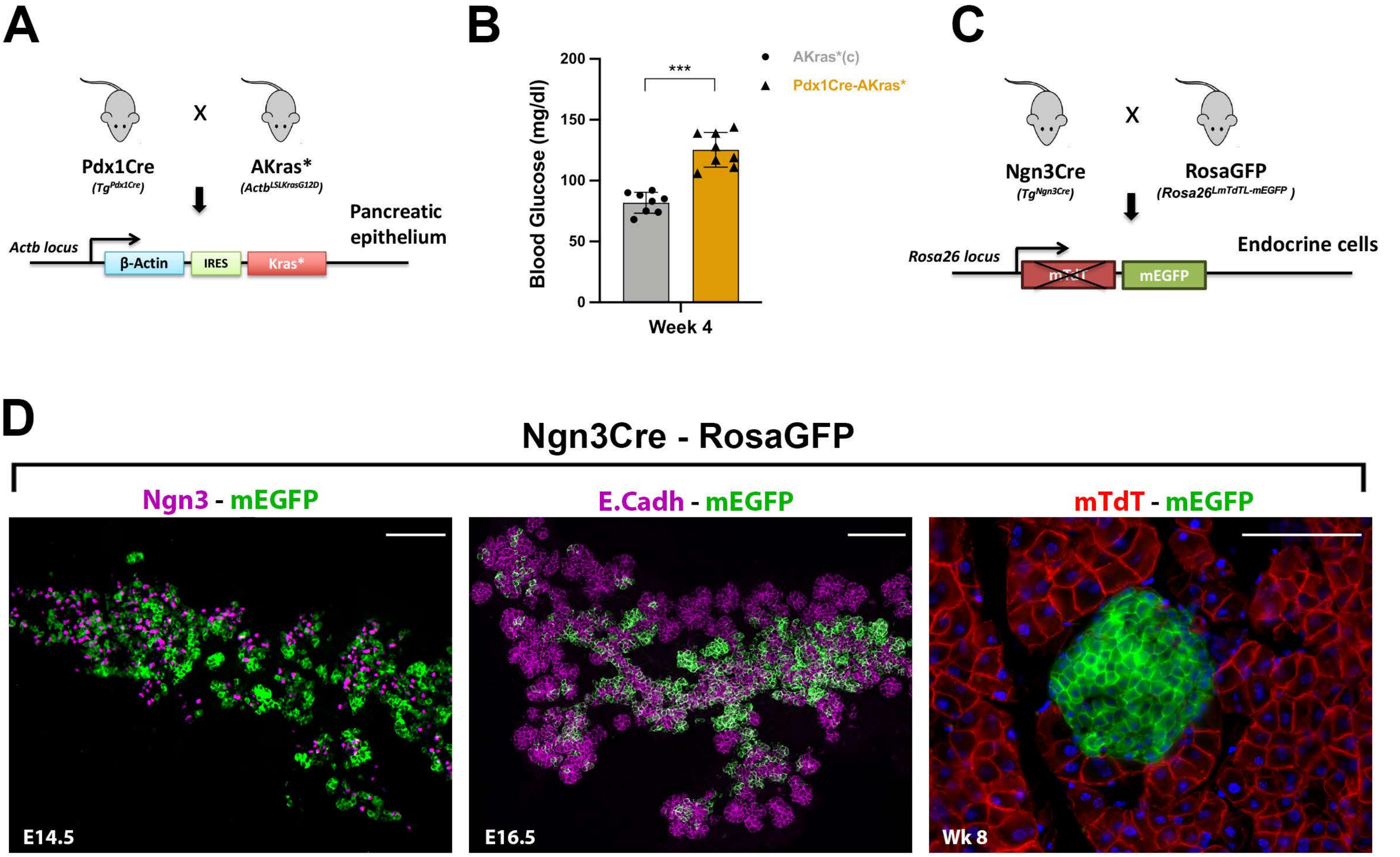

Figure S1

Supplemental Figure 2

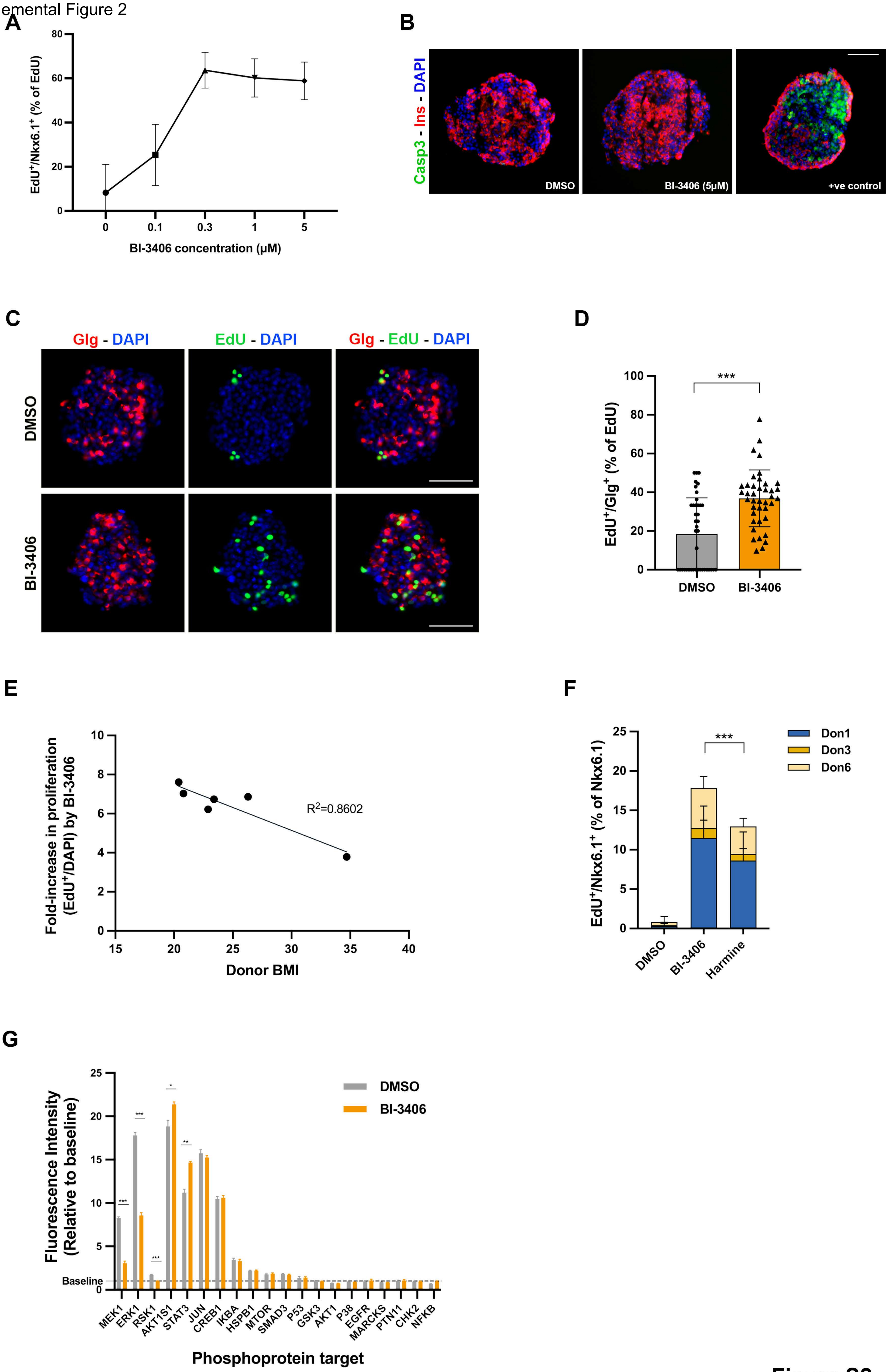

Figure S2

Supplemental Figure 3

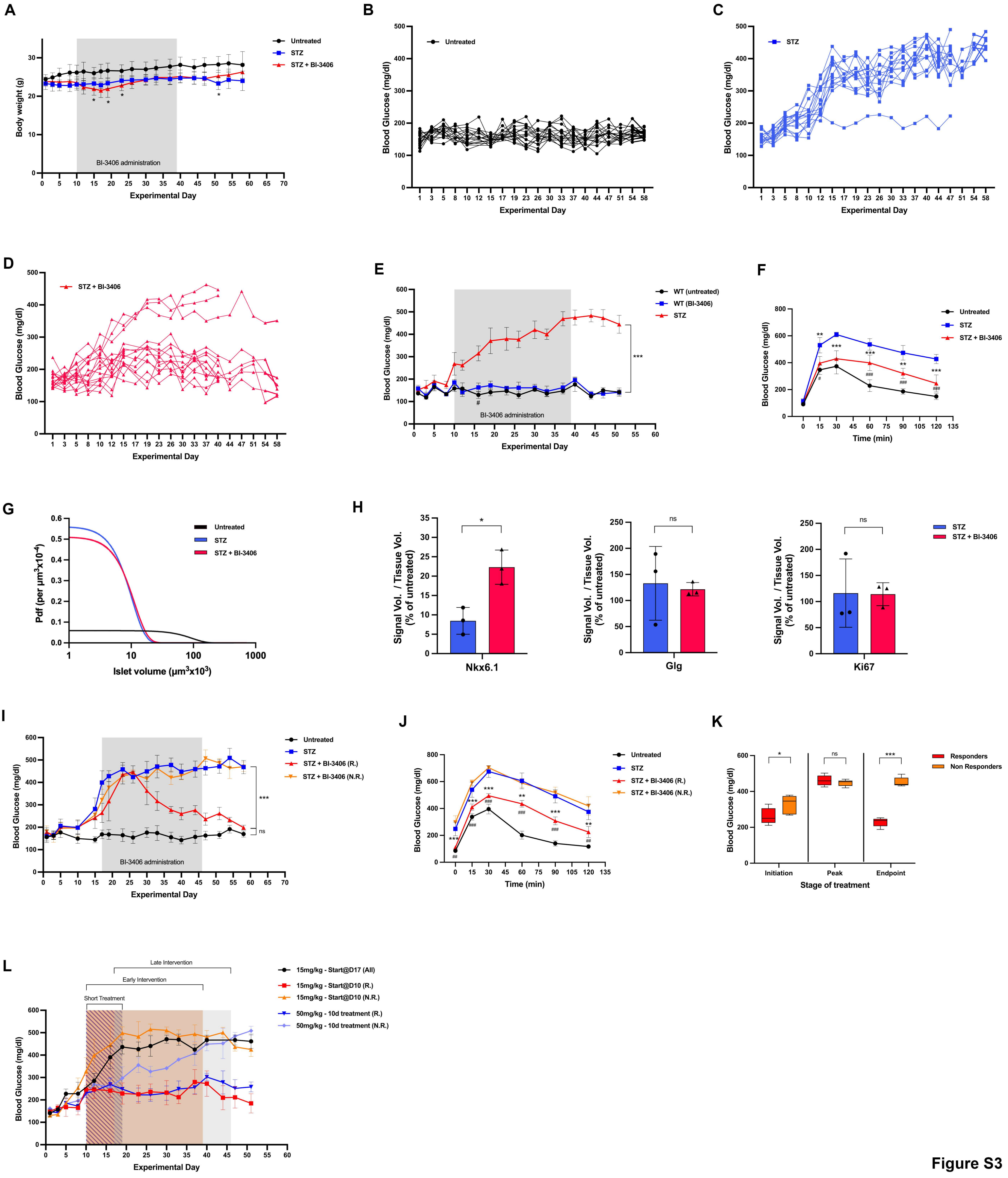

Figure S3

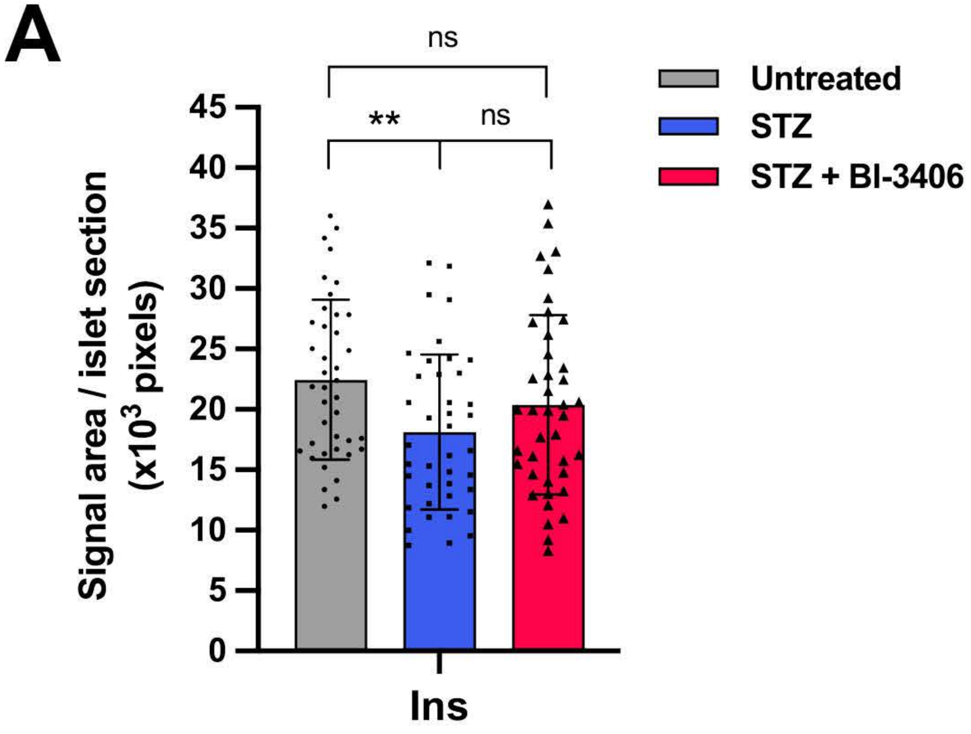
A
ns
**
ns
45
40
35
30
25
20
15
10
5
0
Signal area / islet section (x10^3 pixels)
Ins
Untreated
STZ
STZ + BI-3406

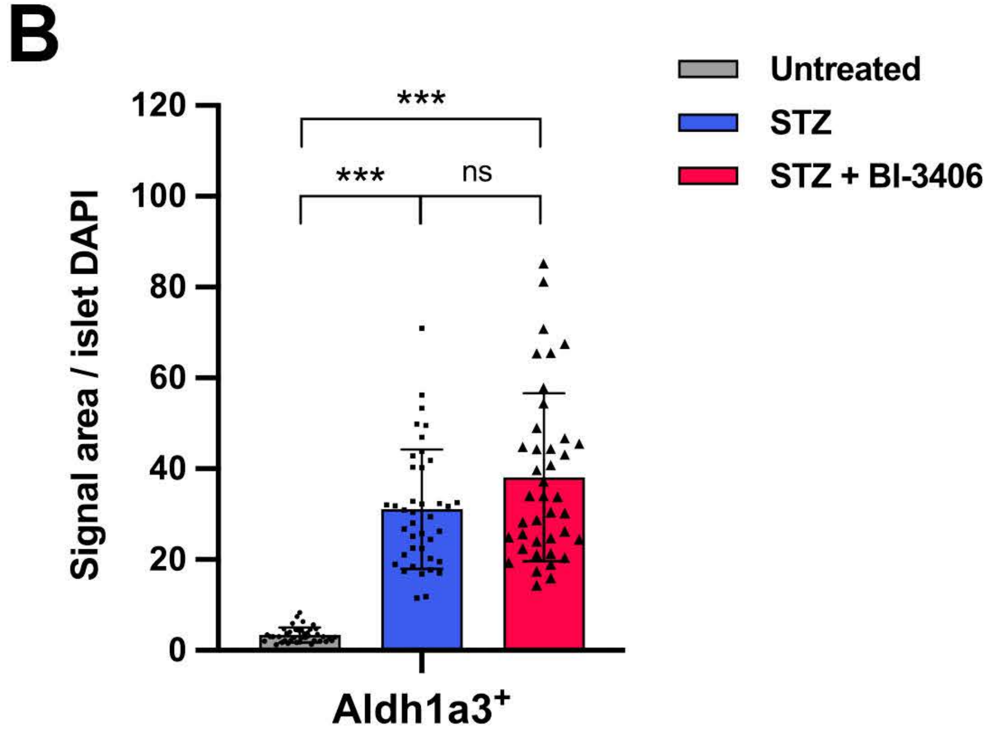
B
***
***
ns
120
100
80
60
40
20
0
Signal area / islet DAPI
Aldh1a3+
Untreated
STZ
STZ + BI-3406

### Figure S4

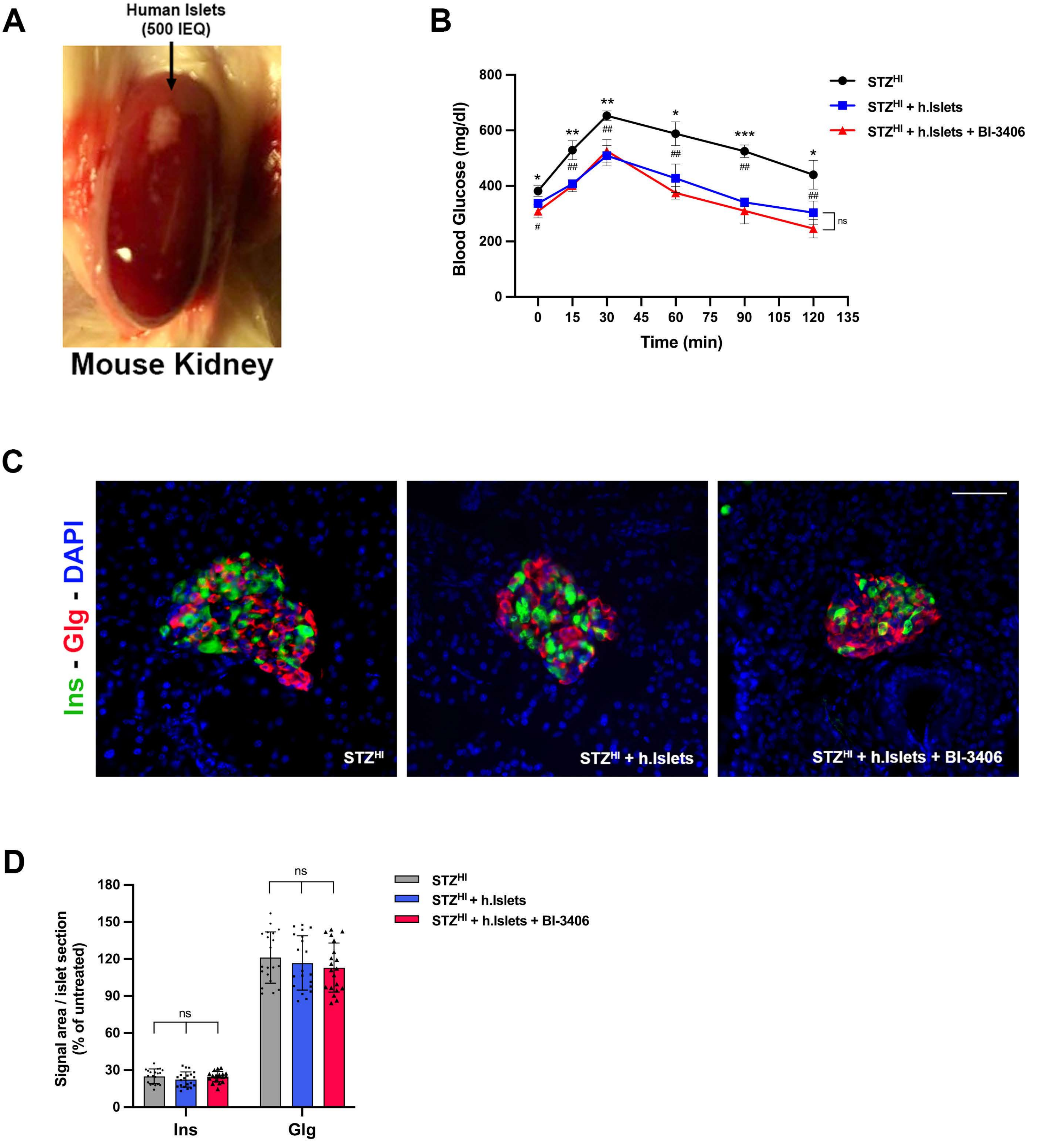
A
Human Islets
(500 IEQ)
Mouse Kidney
B
800
600
400
200
0
Blood Glucose (mg/dl)
0 15 30 45 60 75 90 105 120 135
Time (min)
STZ^HI
STZ^HI + h.Islets
STZ^HI + h.Islets + BI-3406
ns
C
Ins - Glg - DAPI
STZ^HI
STZ^HI + h.Islets
STZ^HI + h.Islets + BI-3406
D
Signal area / islet section
(% of untreated)
180
150
120
90
60
30
0
ns
ns
Ins
Glg
STZ^HI
STZ^HI + h.Islets
STZ^HI + h.Islets + BI-3406

Figure S5
